# Normalization and gene selection for single-cell RNA-seq UMI data using sampling-adjusted sums of squares of Pearson residuals with a Poisson model

**DOI:** 10.1101/2023.12.21.572783

**Authors:** Victor Klebanoff

**Affiliations:** none

**Keywords:** Gene expression, Single-cell RNA-seq, Normalization, Variable genes, Feature selection, Pearson residuals

## Abstract

***SCTransform*** in ***Seurat*** and ***scanpy***.***experimental***.***pp***.***recipe pearson residuals*** (***scanpy*** henceforth) normalize UMI counts as Pearson residuals of negative binomial models. Residual variance scores genes for downstream analysis. Although we observed that both methods usually assign the highest scores to the same genes, for many highly ranked genes (e.g. among the top 2,000) scores may be unstable – not robust to the selection of cells used to calculate residuals. As an alternative, we consider the Poisson model, for which a natural score is the mean sum of squares of Pearson residuals. We show that these scores can be unstable if a gene’s nonzero UMI counts are concentrated on a small number of cells. This explains the instability for ***scanpy*** because of its similarity to the Poisson model. We define a metric for genes’ instability and observe that for all three methods it is negatively correlated with the number of cells on which genes’ counts are nonzero. To reduce the instability of scores based on the Poisson model, we score each gene using multiple random samples of approximately half of the cells. The minimum of these values defines a “sampling-adjusted” score. For data that we analyzed, these are more stable than scores from ***SCTransform*** and ***scanpy*** while generally agreeing with them on the highest ranked genes. As a second criterion to compare our proposal with ***SCTransform***, we use differential expression analysis. For genes with high scores, the residuals’ Kruskal-Wallis H-statistics are generally greater for our method than for ***SCTransform*** and are more highly correlated with our method’s scores.

## 1 Introduction

Numerous software tools are available to analyze single-cell RNA-seq (scRNA-seq) data quantified as UMI (unique molecular identifier) counts. Luecken and Theis (2019) and Amezquita et al. (2020), survey solutions for tasks including normalization, feature selection, dimensionality reduction, clustering, and analysis of differential expression.

Idiosyncratic proposals include using unnormalized counts (Church et al., 2022) or binarized counts (Qiu, 2020). Several approaches normalize counts by cell sequencing depth, add a pseudo-count, often equal to 1, then log-transform the result. Lun (2018) identifies situations when pseudo-counts other than 1 are preferable. Ahlmann-Eltze and Huber (2021) consider alternative transformations and Booeshaghi et al. (2022) recommend a heuristic combining proportional fitting with log-transformation; both emphasize the importance of monotone transformations for downstream data interpretation.

Alternatively, Townes et al. (2019) avoid log-transformation. They propose selecting genes with large deviance and use generalized principal component analysis to reduce dimensions. Similarly, Hafemeister and Satija (2019), with ***SCTransform***, and Lause et al. (2021), with ***scanpy***, avoid log-transformation by normalizing UMI counts as Pearson residuals of negative-binomial models; analysis is performed with the residuals.

There appears to be no consensus on methodologies for evaluating analytical approaches. Church et al. (2022) and Townes et al. (2019) focus on clustering accuracy, applying their methods to benchmark data sets, for which cells’ ground truth classes are given. For two data sets associated with published results, Qiu (2020) compares clusters obtained with their algorithm to those found by ***Seurat*** (Hao et al. (2021)). Identifying highly variable genes is an objective for both Hafemeister and Satija (2019) and Lause et al. (2021). For the 33k PBMC data set investigated by both authors, Hafemeister and Satija (2019) report that the highest-ranked genes found by their approach are exclusively cell-type markers. They contrast their method with log-normalization by comparing UMAP embeddings and the results of differential expression analysis.

We propose a modification of the approaches of ***SCTransform*** and ***scanpy***, using a Poisson model to calculate Pearson residuals. To reduce the influence of cells that can make disproportionate contributions to a gene’s score, we calculate provisional scores for a gene as the mean sum of squares of the residuals using random samples of approximately half of the cells. The minimum of the provisional scores defines the gene’s sampling-adjusted score. This identifies genes for which the sum of squares of residuals taken over all cells is large, but at least one value calculated with a sample of cells is small. Since the characterization of these genes’ variability is not robust to the set of cells used for calculation, their data may be unreliable for downstream analysis. Such genes are typically assigned low scores.

We show how genes with nonzero counts on a small number of cells or, alternatively, many nonzero counts, but only a few large ones, can have unstable provisional scores. The residual-clipping heuristic used in ***SCTransform*** and ***scanpy*** identifies some of these.

For each of the five publicly available data sets that we analyzed, the same genes are generally assigned high scores by all three methods: ***SCTransform, scanpy***, and ours, which we call ***nru*** (**n**ormalize and **r**ank genes using **U**MI counts).

Our approach to comparing these methods is influenced by the discussion in Yip et al. (2019) of the sensitivity of results to samples of cells used for analysis. Similar considerations arise in analysis of the stability of clusterings, as in Hennig (2007), Lun (2019), Peyvandipour et al. (2020), and Patterson-Cross et al. (2021).

To analyze the stability of scores calculated by ***SCTransform, scanpy***, and ***nru***, we select a random sample of approximately half of the cells in the count matrix. The cell sample defines a count sub-matrix. Each method is applied to the sub-matrix and its complement. The confusion matrix comparing the ranks of genes’ scores for the two samples provides one characterization of stability. Additionally, we define a measure of instability for genes’ scores and compare its distribution for ***nru*** with ***scanpy*** and ***SCTransform***. We find that ***nru*** ‘s scores are more stable and that the instability of genes’ scores is negatively correlated with the number of cells in which their counts are nonzero.

We perform differential expression analysis for genes assigned high scores by ***SCTransform*** and ***nru*** by calculating Kruskal-Wallis H-statistics for the Pearson residuals using clusterings (cell types) provided with the count data sets. For genes with high scores, the H-statistics are generally more highly correlated with ***nru*** ‘s scores than ***SCTransform*** ‘s. Similarly, for subsets of the genes scored highest by each method, the distributions of H-statistics are generally larger for genes with high ***nru*** scores.

Finally, we obtain what may be new insights from a closed-form expression for the mean sum of squares of Pearson residuals of a Poisson model: it depends only on the cells in which a gene’s counts are nonzero. A cell’s contribution depends only on its sequencing depth, the gene’s count in the cell, and the UMI matrix’s total count and number of cells. In particular, the formula displays the role of sequencing depth in characterizing genes’ variability.

Our programs, in Jupyter notebooks containing the results in this paper and Supplement 1, are available at https://github.com/victorkleb/scRNA-seq_Pr.

## 2 Methods

We initially quantify the variability of gene *g* as the mean sum of squares of its Pearson residuals calculated with a Poisson model, denoted ***M*** _*g*_. We derive a formula for ***M*** _*g*_ which shows that it depends only on data for cells in which the gene has nonzero counts (we refer to these as “nonzero cells”) – and on the number of cells in the UMI matrix and its total count. The formula explains why ***M*** _*g*_ can be large for genes with few nonzero cells. Similarly, it shows how ***M*** _*g*_ can be large if the gene *g* has large UMI counts in few cells and its remaining nonzero counts are small. For these genes, ***M*** _*g*_ may not be robust to the influence of a small number of cells, hence the gene’s data may be unreliable. If a UMI count matrix contains sufficient data to permit valid analysis, independent estimates of ***M*** _*g*_ calculated with random samples of half of the cells should be consistent.

We randomly draw *K* samples of approximately half of the cells, denoting a sample as *S*_*k*_. We calculate ***M*** _*g*_(*S*_*k*_) for the UMI count sub-matrix containing the cells in sample *S*_*k*_, then define the sampling-adjusted mean sum of squares of Pearson residual, denoted ***A***_*g*_, as the minimum of these *K* estimates: ***A***_*g*_ = min *{****M*** _*g*_(*S*_1_), ***M*** _*g*_(*S*_2_), *· · ·* , ***M*** _*g*_(*S*_*K*_)*}*. We use ***A***_*g*_ to score each gene.

### 2.1 Notation

Consider a UMI count matrix with *X*_*gc*_ representing the count for gene *g* in cell *c*. Let

*G*_*g*_ = Σ_*c*_ *X*_*gc*_ the total count for gene *g*

*π*_*c*|*g*_ = *X*_*gc*_*/G*_*g*_ the fraction of the total count of gene *g* that is in cell *c*

*D*_*c*_ =Σ_*g*_ *X*_*gc*_ the total count for cell *c* – its sequencing depth

*T* = Σ_*gc*_ *X*_*gc*_ the total of all counts in the matrix

*C* = the total number of cells in the matrix

*D* = *T/C* the mean sequencing depth of the matrix

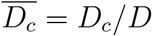 the normalized sequencing depth of cell *c*

### 2.2 A Poisson model for UMI count data

We assume a Poisson model for the UMI counts. As Lause et al. (2021) observe, the maximum likelihood estimate of the count equals 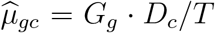 and the Pearson residual for cell *c* and gene *g* equals 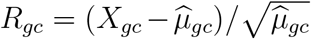 . We?quantify the variability of gene *g* as the mean sum of squares of its Pearson residuals 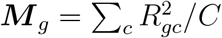.

### 2.3 A closed-form expression for *M* _*g*_

In Appendix A we show that

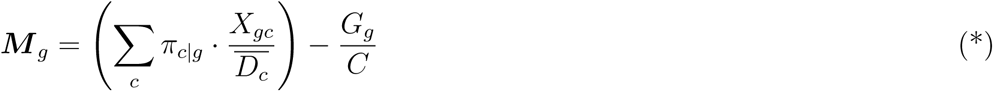

We make the following observations:

- The summation includes *only* cells in which gene *g* has nonzero counts. We denote the number of these cells by ***n***_*g*_.
- The mean value of the normalized sequencing depths 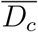 over all cells in the count matrix equals 1.
- The subtracted term *G*_*g*_*/C* is negligible for lowly expressed genes.
- Since ∑ _*c*_ *π*_*c*|*g*_ = 1, the summation is a convex combination of the terms 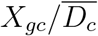 – each is the gene’s count in cell *c* divided by the cell’s normalized sequencing depth.

The formula for ***M*** _*g*_ (*) explains the large values observed for some genes with a nonzero UMI count in only one cell. In this case, the summation reduces to a single term, for – for convenience – cell 1. The gene’s total count equals its count in cell 1: *G*_*g*_ = *X*_*g*1_ and *π*_1|*g*_ = 1, making 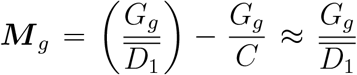, because *G*_*g*_ *≪ C*. That is, ***M*** _*g*_ is approximately the gene’s count divided by the normalized sequencing depth of its nonzero cell. If the cell’s sequencing depth is below average, ***M*** _*g*_ can be large. In the data sets that we studied, *G*_*g*_ equals 1 for over 95% of genes with only one nonzero cell. For these genes 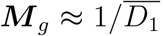. If *G*_*g*_ *>* 1, then ***M*** _*g*_ is even larger.

This also explains large values of ***M*** _*g*_ for some genes with nonzero UMI counts in a small number of cells. ***M*** _*g*_ is the convex combination of ratios of the form just described.

### 2.4 Genes with unreliable data: a toy example

Analysis of the impact of gene clipping in ***scanpy*** shows that ***M*** _*g*_ can be very large when a gene has large counts in one, or few, cells and small nonzero counts otherwise. Despite the large ***M*** _*g*_ (or, in ***scanpy***, residual variance) the gene’s data should be excluded from downstream analysis if the gene cannot be reliably characterized as highly variable: were a few cells absent from the sample, the gene would be nondescript. We believe that this is the underlying problem that clipping is meant to solve. To motivate the use of random samples of cells to identify these genes, we give a toy example. Because the subtracted term *G*_*g*_*/C* in (*) is negligible for lowly expressed genes, we ignore it and consider 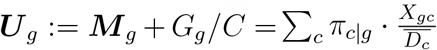

Suppose

- for simplicity, that the normalized sequencing depth 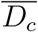 is 1 for every cell, making ***U*** _*g*_ = ∑_*c*_ *π*_*c*|*g*_ *· X*_*gc*_
- and that *N* cells have nonzero counts: for cell 1, the count *X*_*g*1_ equals *B*; for the remaining *N −* 1 cells, all counts are 1.

Hence

- the gene’s total count *G*_*g*_ = *B* + *N −* 1.
- and 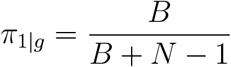 with 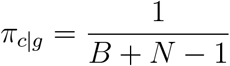 for the remaining nonzero cells.

Then 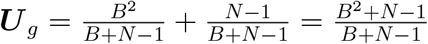. A simple example shows how this can be large in a case of no biological interest. Suppose that *N* = 100 and *B* = 25 (100 nonzero cells; 1 count equals 25; the other 99 counts equal 1). Then 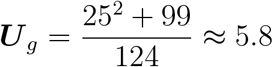. Cell 1 contributes 5.0 – 87% of the total. In the 33k PBMC data set, the 99^*th*^ percentile of the distribution of ***M*** _*g*_ is 4.7. The value from this toy example, 5.8, would be among the largest 1%. To show that this is unreliable, split the set of cells into two subsets and suppose that each includes 50 nonzero cells. Calculate ***M*** _*g*_ using each subset: for the subset containing cell 1, ***M*** _*g*_ = 9.1; for the complementary subset ***M*** _*g*_ = 1.0.

### 2.5 *A*_*g*_ – the sampling-adjusted mean sum of squares of Pearson residuals

The example above shows how complementary samples of cells can give different estimates of ***M*** _*g*_ if only one cell has a large nonzero count. For the general case, when a small number of cells have large counts, we propose selecting several random samples of the count matrix, each containing approximately half the cells, calculating ***M*** _*g*_ with each, then quantifying the gene’s variability as the minimum of these values. We apply the following algorithm using 40 samples:

for *k in {*1, *· · ·* , 20*}*

select *S*_*k*_, a random sample of approximately half of the cells

calculate ***M*** _*g*_(*S*_*k*_) for each gene using counts for the cells in *S*_*k*_

calculate 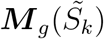 for each gene using counts for the cells in 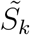 , the complement of *S*_*k*_ assign 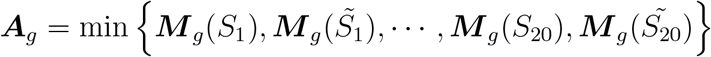

## 3 Results and discussion

### 3.1 Data sets

We studied five publicly available data sets (Table 1).

**Table 1.**
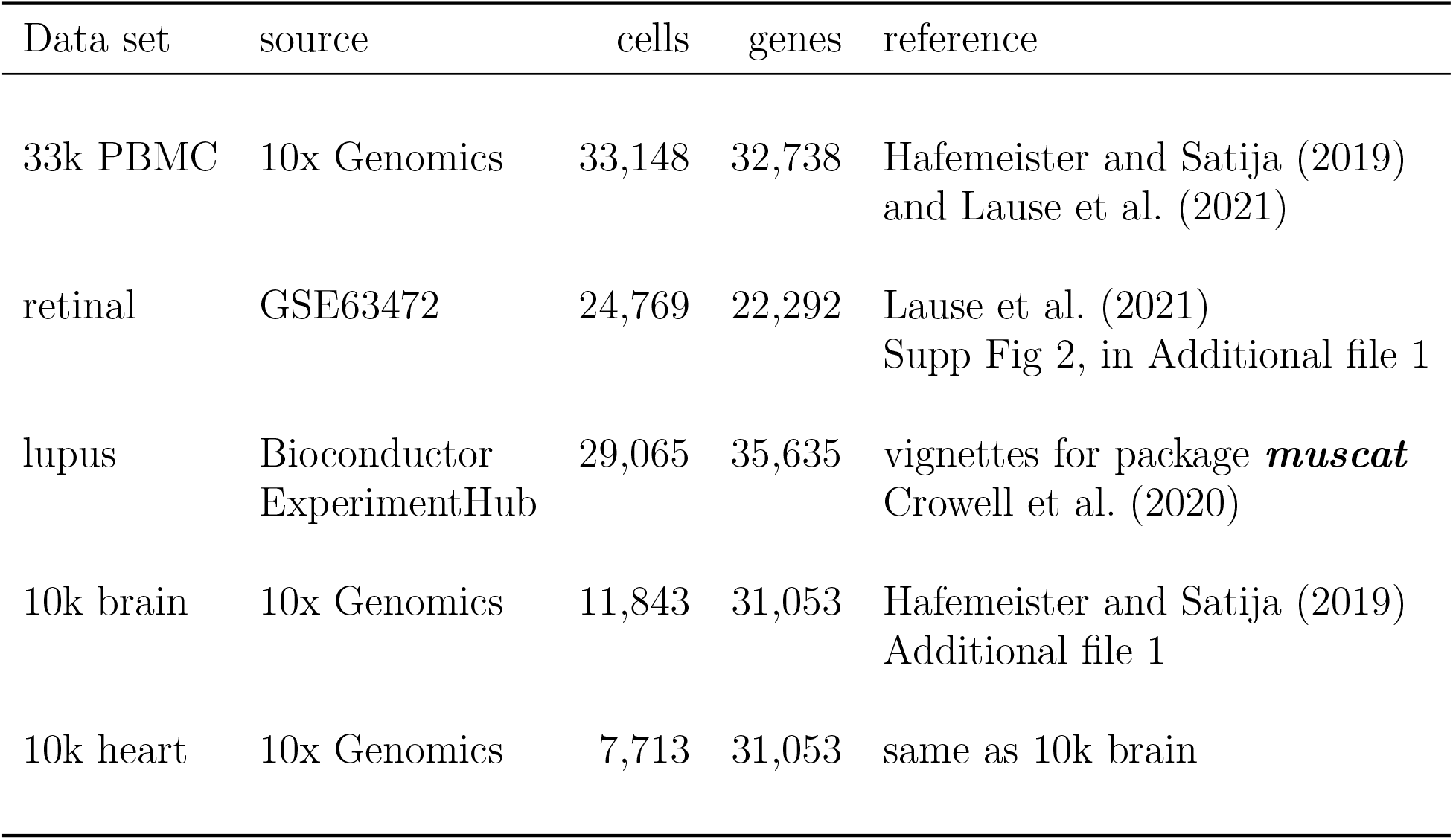
Data sets used. The GSE63472 data set includes counts for 24,658 genes in 49,300 cells. We used the same subset as Lause et al. (2021)

| Data set | source | cells | genes | reference |
| --- | --- | --- | --- | --- |
| 33k PBMC | 10x Genomics | 33,148 | 32,738 | Hafemeister and Satija (2019)<br>and Lause et al. (2021) |
| retinal | GSE63472 | 24,769 | 22,292 | Lause et al. (2021)<br>Supp Fig 2, in Additional file 1 |
| lupus | Bioconductor<br>ExperimentHub | 29,065 | 35,635 | vignettes for package *muscat*<br>Crowell et al. (2020) |
| 10k brain | 10x Genomics | 11,843 | 31,053 | Hafemeister and Satija (2019)<br>Additional file 1 |
| 10k heart | 10x Genomics | 7,713 | 31,053 | same as 10k brain |

This paper gives results for the 33k PBMC data set. The data, available online from 10x Genomics, are associated with the paper by Zheng et al. (2017). The retinal data were studied by Lause et al. (2021); results are in the supplementary material (Figure 2) to their paper. The lupus data are used in two Bioconductor vignettes for ***muscat*** (Crowell et al., 2020). Results for the heart and brain data are reported in Additional File 1, “Application of ***sctransform*** to additional UMI data sets” with Hafemeister and Satija (2019). Our results for the retinal, lupus, heart, and brain data sets are summarized in Supplement 1.

### 3.2 Performance of *nru*

Figure 1A compares ***M*** _*g*_ to ***n***_*g*_ for all 20,678 genes with nonzero counts. (Points represent genes in all scatter plots.) It suggests that many genes with large values of ***M*** _*g*_ have few nonzero cells. Table 2 confirms this. ***M*** _*g*_ is greater than 2 for 1,266 genes. Of these, 334 have 50 or fewer nonzero cells.

**Table 2.** 33k PBMC data: genes with ***n***_*g*_ *≤* 10 are responsible for 210 of the 1,266 values of ***M*** _*g*_ *>* 2; genes with 11 *≤* ***n***_*g*_ *≤* 50 are responsible for an additional 124.

| $\mathbf{M}_g$ | $\leq 1$ | $1 < -2$ | $> 2$ | Total |
| --- | --- | --- | --- | --- |
| $\mathbf{n}_g$ | | | | |
| 1-10 | 2,768 | 2,492 | 210 | 5,470 |
| 11-50 | 1,069 | 1,723 | 124 | 2,916 |
| 51-100 | 334 | 839 | 94 | 1,267 |
| 101-1,000 | 802 | 4,826 | 507 | 6,135 |
| 1,001-10,000 | 55 | 4,169 | 190 | 4,414 |
| 10,001+ | 0 | 335 | 141 | 476 |
| Total | 5,028 | 14,384 | 1,266 | 20,678 |

**Figure 1.**
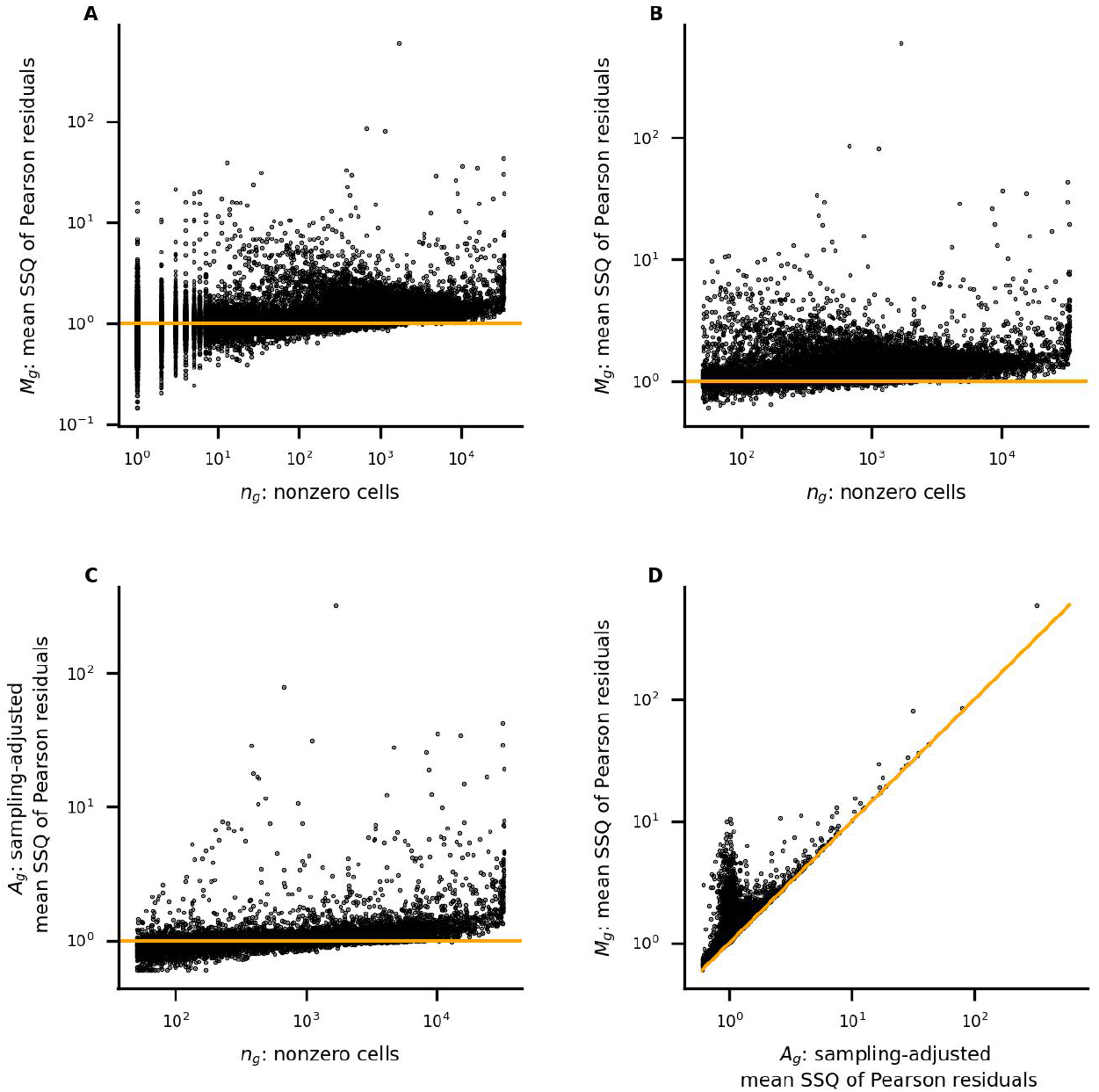
33k PBMC data: (**A**) the mean SSQ of Pearson residuals is large for many genes with few nonzero cells, for which data are unlikely to be reliable for downstream analysis; (**B**) restricting to genes with at least 50 nonzero cells retains 12,324; (**C**) ***A***_*g*_ is smaller than ***M*** _*g*_ for many genes with few nonzero cells; (**D**) adjustment reduces some scores by an order of magnitude

It would be appealing to identify a threshold *θ* such that data for few, if any, genes with ***n***_*g*_ *< θ* are reliable for downstream analysis. Since we could not do this, we arbitrarily exclude all genes with fewer than 50 nonzero cells. For the 33k PBMC data set, this retains 12,324 genes. Figure 1B shows the relation between ***M*** _*g*_ and ***n***_*g*_ for them.

The impact of applying the algorithm of section 2.5 is illustrated in Figures 1C and 1D. Compared with 1B, 1C shows that ***A***_*g*_ is much smaller than ***M*** _*g*_ for many genes, particularly those with fewer nonzero cells; 1D compares ***M*** _*g*_ with ***A***_*g*_. We propose excluding from downstream analysis most genes represented by points above the diagonal in the neighborhood of (1,1).

To evaluate the stability of ***A***_*g*_ we select *S*, a random sample of approximately half the cells, calculate ***A***_*g*_(*S*) for each gene *g* restricting to cells in *S*, then calculate 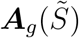 restricting to cells in 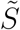, the complement of *S*.

Table 3 displays the confusion matrix comparing ranks of ***A***_*g*_(*S*) with 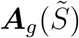: 19 genes are ranked in the top 20 in both samples; 47 in the top 50; 99 in the top 100; 189 in the top 200; 446 in the top 500. Alternatively, 99% of all genes ranked in the top 500 for each sample are in the top 2,000 for the other. Restricting to genes with 50 or more nonzero cells in both *S* and 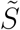 retains the 10,908 represented in the table.

**Table 3.** 33k PBMC data: comparing ranks of ***A***_*g*_ calculated with *S* and 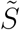.

| sample $\tilde{S}$ | 1-20 | 21-50 | 51-100 | 101-200 | 201-500 | 501-2000 | 2001+ | Total |
| --- | --- | --- | --- | --- | --- | --- | --- | --- |
| sample $S$ | | | | | | | | |
| 1-20 | 19 | 1 | 0 | 0 | 0 | 0 | 0 | 20 |
| 21-50 | 1 | 26 | 3 | 0 | 0 | 0 | 0 | 30 |
| 51-100 | 0 | 3 | 46 | 0 | 1 | 0 | 0 | 50 |
| 101-200 | 0 | 0 | 1 | 89 | 10 | 0 | 0 | 100 |
| 201-500 | 0 | 0 | 0 | 11 | 235 | 50 | 4 | 300 |
| 501-2000 | 0 | 0 | 0 | 0 | 49 | 980 | 471 | 1,500 |
| 2001+ | 0 | 0 | 0 | 0 | 5 | 470 | 8,433 | 8,908 |
| Total | 20 | 30 | 50 | 100 | 300 | 1,500 | 8,908 | 10,908 |

For gene *g*, the figure of merit

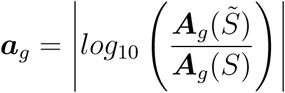

quantifies the instability of ***A***_*g*_ calculated with *S* and 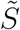.

The scores ***A***_*g*_(*S*) and 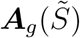 are compared in Figure 2A. In Figure 2B, the instability ***a***_*g*_ is compared to ***A***_*g*_ (calculated with all cells). Scores agree within a factor of 2 for all genes except IGLL5. Small values of ***a***_*g*_ for large values of ***A***_*g*_ correspond to the large values in Figure 2A near the diagonal. Figure 2C compares ***a***_*g*_ to ***n***_*g*_ for 1,932 genes satisfying two conditions: they are among 10,908 for which ***a***_*g*_ was calculated and their values of ***A***_*g*_ are among the largest 2,000. ***a***_*g*_ tends to be smaller for large values of ***n***_*g*_. The Spearman correlation between ***a***_*g*_ and ***n***_*g*_ equals -0.17.

**Figure 2.**
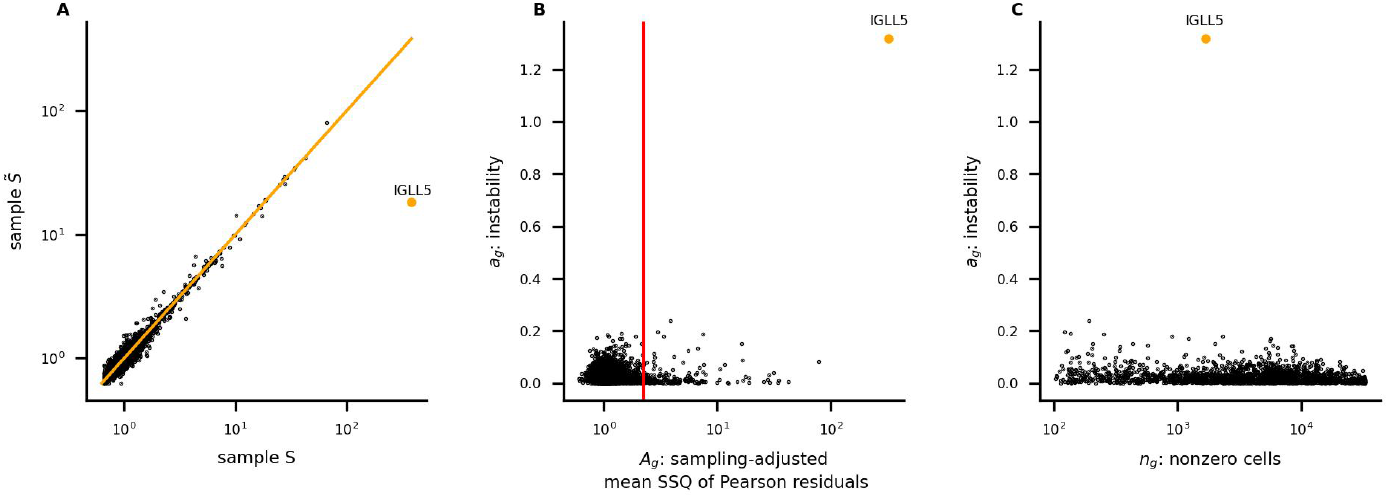
33k PBMC data: (**A**) large values of ***A***_*g*_(*S*) and 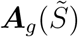 agree closely, except for the outlier IGLL5; (**B**) the red vertical line marks the 200^*th*^ ranked value of ***A***_*g*_ calculated with all cells; (**C**) for the 1,932 genes with the largest values of ***A***_*g*_, for which both ***A***_*g*_(*S*) and 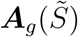 were calculated, instability tends to be larger for genes with fewer nonzero cells (Spearman correlation: -0.17)

Despite the disparity between the scores ***A***_*IGLL*5_(*S*) and 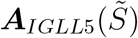, the situation seems benign because both are large. Note that IGLL5 is highly ranked by all three methods: 6^*th*^ highest by ***SCTransform***, 14^*th*^ by ***scanpy***, and first by ***nru***. In the 10k heart data set, similar behavior for three genes may reflect a quality control issue – counts concentrated on a small number of cells. Indeed, this may be a consideration for IGLL5: it has 1,679 nonzero cells and its total count equals 32,897 (its total count per nonzero cell is the 11^*th*^ highest in the data set). Over 10% of its total count, 3,866, is in one cell; six cells are responsible for more than 50% of its total count and ten cells are responsible for more than 70%.

### 3.3 Relation to *scanpy*

Code was excerpted from lines 26-64 of the function _*pearson_residuals* in _*normalization*.*py* on the github page https://github.com/scverse/scanpy/tree/master/scanpy/experimental/pp

***Scanpy*** and ***nru*** assign the highest scores to many of the same genes. ***Scanpy*** follows ***SC-Transform*** by clipping residuals, which identifies some genes to exclude from downstream analysis.

Figure 3A compares ***M*** _*g*_ with scores calculated by ***scanpy*** *without* clipping. Their Spearman correlation is 1.0. The effect of the negative binomial model is small: for 99% of genes, the difference is less than 3%, for 97% of genes, it is less than 1%.

**Figure 3.**
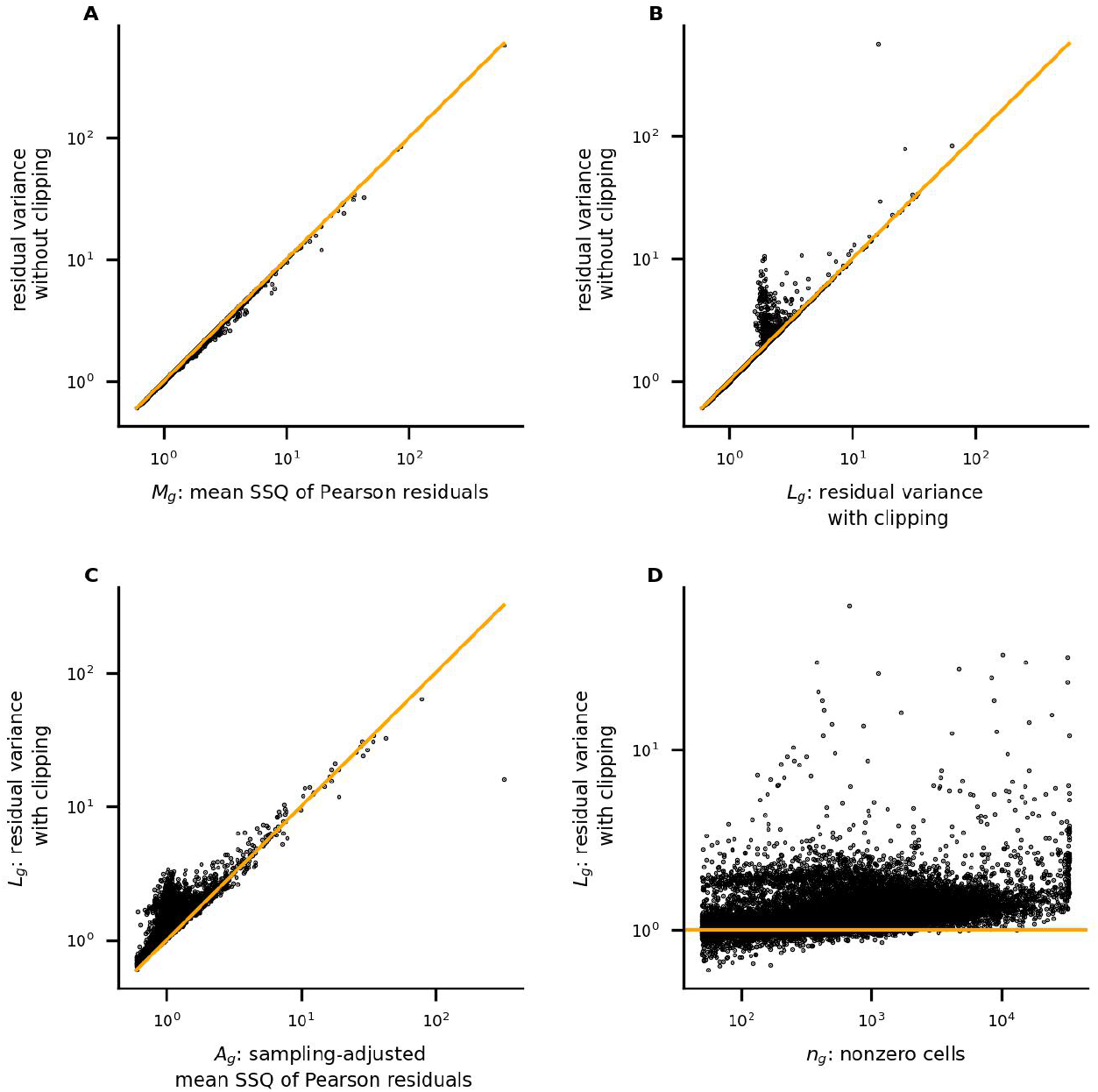
33k PBMC data: (**A**) scores calculated by ***scanpy*** without clipping agree closely with ***M*** _*g*_; the effect of the negative binomial model is small; (**B**) clipping affects scores for 413 genes; (**C**) ***L***_*g*_ and ***A***_*g*_ are large for many of the same genes; (**D**) ***scanpy*** assigns high scores to many genes with fewer than 1,000 nonzero cells

Figure 3B shows the effect of clipping. We denote the ***scanpy*** score, with clipping, by ***L***_*g*_. Figure 3C compares ***L***_*g*_ with ***A***_*g*_. They are large for many of the same genes. Even with clipping, ***scanpy*** assigns higher scores to the genes represented by points above the diagonal in the neighborhood of (1,1). For many of these genes, provisional scores (used to calculate ***A***_*g*_) vary by a factor of two or more; their data may be unreliable for downstream analysis. In Figure 3D, ***L***_*g*_ is compared to the number of nonzero cells ***n***_*g*_. Comparison with Figure 1C shows that ***L***_*g*_ is more likely than ***A***_*g*_ to be large for genes with fewer than 1,000 nonzero cells.

Table 4 displays the confusion matrix comparing ranks of ***L***_*g*_ with ***A***_*g*_: 19 genes are ranked in the top 20 by both; 44 in the top 50; 89 in the top 100.

**Table 4.**
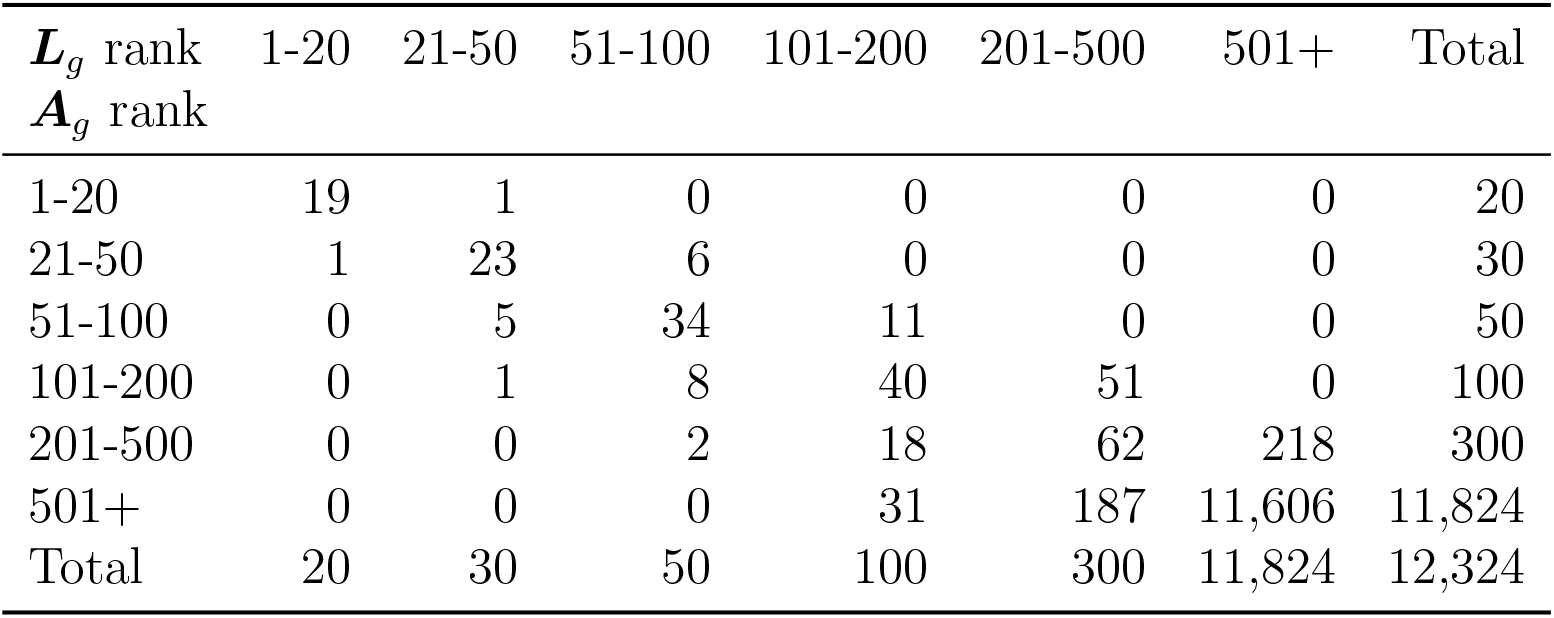
33k PBMC data: comparing ranks of ***L***_*g*_ with ***A***_*g*_.

| $\mathbf{L}_g$ rank<br>$\mathbf{A}_g$ rank | 1-20 | 21-50 | 51-100 | 101-200 | 201-500 | 501+ | Total |
| --- | --- | --- | --- | --- | --- | --- | --- |
| 1-20 | 19 | 1 | 0 | 0 | 0 | 0 | 20 |
| 21-50 | 1 | 23 | 6 | 0 | 0 | 0 | 30 |
| 51-100 | 0 | 5 | 34 | 11 | 0 | 0 | 50 |
| 101-200 | 0 | 1 | 8 | 40 | 51 | 0 | 100 |
| 201-500 | 0 | 0 | 2 | 18 | 62 | 218 | 300 |
| 501+ | 0 | 0 | 0 | 31 | 187 | 11,606 | 11,824 |
| Total | 20 | 30 | 50 | 100 | 300 | 11,824 | 12,324 |

The stability of ***L***_*g*_ is evaluated for the same samples *S* and 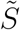 as ***A***_*g*_. The confusion matrix comparing ranks of ***L***_*g*_(*S*) with 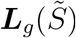 is displayed in Table 5. Results for ***L***_*g*_ are less stable than for ***A***_*g*_ (Table 3): 19 genes are ranked in the top 20 in both samples; 47 in the top 50; 92 in the top 100; 141 in the top 200; 252 in the top 500. For ***A***_*g*_ all genes in the top 200 in one sample are in the top 500 in the other. This is not the case for ***L***_*g*_: 43 genes in the top 200 in *S* are not in the top 500 in 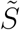; 45 in the top 200 in 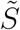 are not in the top 500 in *S*.

**Table 5.** 33k PBMC data: comparing ranks of ***L***_*g*_ calculated with *S* and 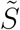.

| sample $\tilde{S}$<br>sample $S$ | 1-20 | 21-50 | 51-100 | 101-200 | 201-500 | 501-2000 | 2001+ | Total |
| --- | --- | --- | --- | --- | --- | --- | --- | --- |
| 1-20 | 19 | 1 | 0 | 0 | 0 | 0 | 0 | 20 |
| 21-50 | 1 | 26 | 3 | 0 | 0 | 0 | 0 | 30 |
| 51-100 | 0 | 3 | 39 | 8 | 0 | 0 | 0 | 50 |
| 101-200 | 0 | 0 | 5 | 36 | 16 | 9 | 34 | 100 |
| 201-500 | 0 | 0 | 1 | 13 | 81 | 61 | 144 | 300 |
| 501-2000 | 0 | 0 | 1 | 7 | 68 | 335 | 1,089 | 1,500 |
| 2001+ | 0 | 0 | 1 | 36 | 135 | 1,095 | 7,641 | 8,908 |
| Total | 20 | 30 | 50 | 100 | 300 | 1,500 | 8,908 | 10,908 |

For gene *g*, instability is calculated as

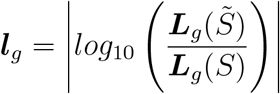

The scores ***L***_*g*_(*S*) and 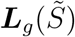 are compared in Figure 4A. In Figure 4B, ***l***_*g*_ is compared to ***L***_*g*_. Figure 4C compares ***l***_*g*_ to ***n***_*g*_ for 1,848 genes satisfying two conditions: they are among 10,908 for which ***l***_*g*_ was calculated and their values of ***L***_*g*_ are among the largest 2,000. The Spearman correlation between ***l***_*g*_ and ***n***_*g*_ equals -0.54.

**Figure 4.**
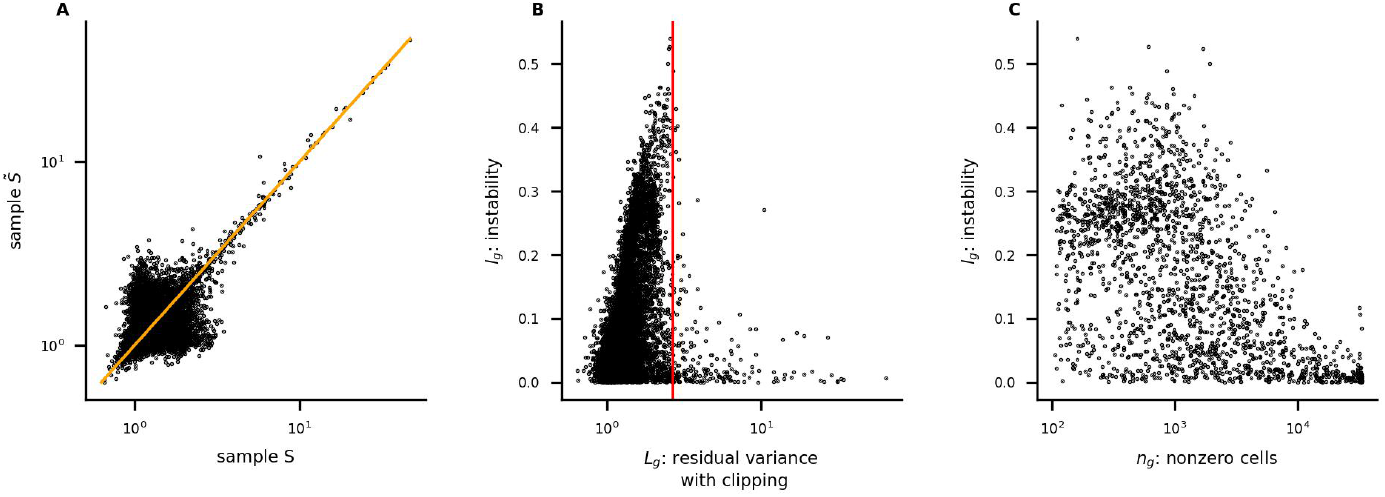
33k PBMC data: (**A**) large values of ***L***_*g*_(*S*) and 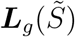 agree closely; (**B**) the red vertical line marks the 200^*th*^ ranked value of ***L***_*g*_ calculated with all cells; (**C**) for the 1,848 genes with the largest values of ***L***_*g*_, for which both ***L***_*g*_(*S*) and 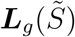 were calculated, instability is larger for genes with fewer nonzero cells (Spearman correlation: -0.54)

In addition to comparing the confusion matrices in Tables 5 and 3, we compared the distributions of ***a***_*g*_ and ***l***_*g*_ (restricting to genes represented in Figures 2C and 4C) by performing Mann-Whitney and Kolmogorov-Smirnov tests.

The Mann-Whitney one-sided test rejects the null hypothesis that ***l***_*g*_ and ***a***_*g*_ have the same distribution in favor of the alternative that the distribution of ***l***_*g*_ is stochastically greater than that of ***a***_*g*_ (p=0). Similarly, the Kolmogorov-Smirnov one-sided test rejects the null hypothesis in favor of the alternative that the CDF of ***l***_*g*_ is less than that of ***a***_*g*_ (p=0). Repeating the analysis with 19 additional pairs of complementary random samples gave the same results.

### 3.4 Relation to *SCTransform*

Pearson residuals and scores were calculated with version 2 of ***SCTransform*** (Choudhary and Satija, 2022). ***S***_*g*_ denotes the score for gene *g*. Figure 5A compares ***S***_*g*_ to ***n***_*g*_. ***S***_*g*_ is very small for most genes with few nonzero cells. Figure 5B compares ***S***_*g*_ with ***A***_*g*_. They are large for many of the same genes.

**Figure 5.**
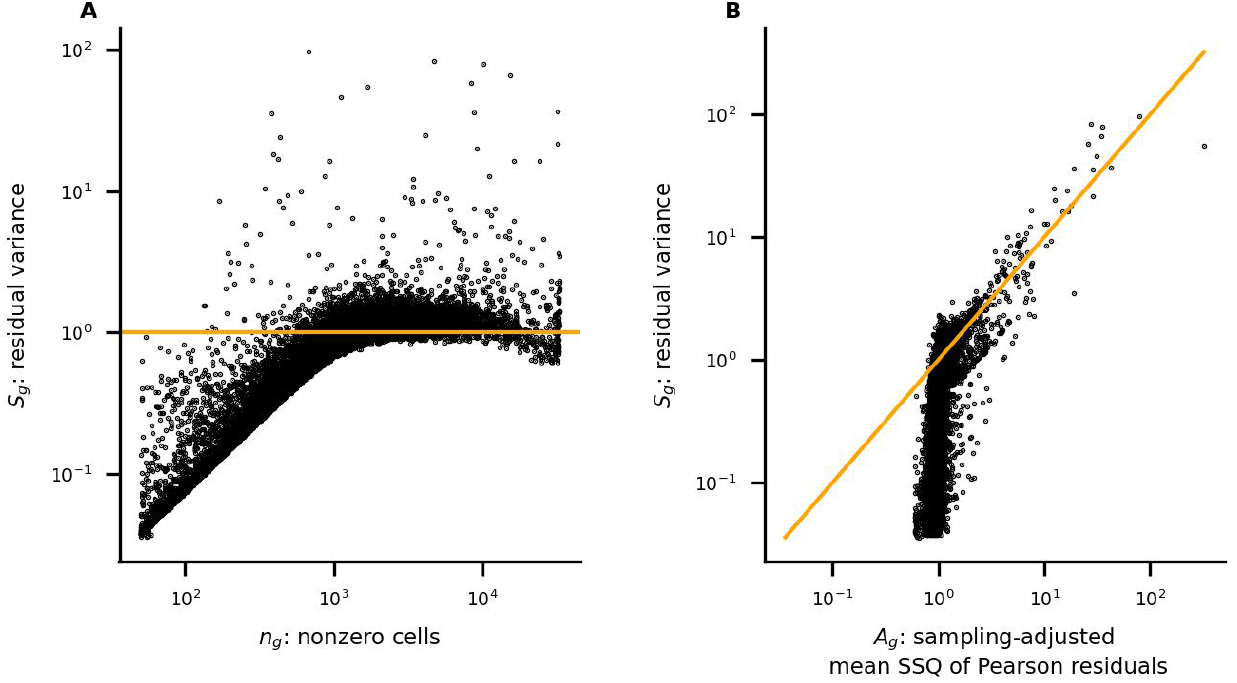
33k PBMC data: (**A**) ***SCTransform*** assigns very low scores to most genes with few nonzero cells; (**B**) ***S***_*g*_ and ***A***_*g*_ are large for many of the same genes

Table 6 displays the confusion matrix comparing ranks of ***S***_*g*_ with ***A***_*g*_: 18 genes are ranked in the top 20 by both; 37 in the top 50; 78 in the top 100.

**Table 6.** 33k PBMC data: comparing ranks of ***S***_*g*_ with ***A***_*g*_.

| $S_g$ rank<br>$A_g$ rank | 1-20 | 21-50 | 51-100 | 101-200 | 201-500 | 501+ | Total |
| --- | --- | --- | --- | --- | --- | --- | --- |
| 1-20 | 18 | 1 | 1 | 0 | 0 | 0 | 20 |
| 21-50 | 2 | 16 | 9 | 3 | 0 | 0 | 30 |
| 51-100 | 0 | 12 | 19 | 5 | 10 | 4 | 50 |
| 101-200 | 0 | 1 | 17 | 22 | 9 | 51 | 100 |
| 201-500 | 0 | 0 | 3 | 52 | 75 | 170 | 300 |
| 501+ | 0 | 0 | 1 | 18 | 206 | 11,599 | 11,824 |
| Total | 20 | 30 | 50 | 100 | 300 | 11,824 | 12,324 |

Hafemeister and Satija (2019) and Lause et al. (2021) debated the variability of the gene MALAT1. It is ranked 11^*th*^ by ***nru*** and 77^*th*^ by ***SCTransform*** (version 2).

The stability of ***S***_*g*_ is evaluated for the same samples *S* and 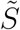 as ***A***_*g*_ and ***L***_*g*_. The confusion matrix comparing ranks of ***S***_*g*_(*S*) with 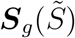 is displayed in Table 7. The top 20 genes in both samples are identical; 49 are in the top 50 in both; 89 in the top 100; 143 in the top 200; 252 in the top 500.

**Table 7.** 33k PBMC data: comparing ranks of ***S***_*g*_ calculated with *S* and 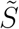.

| sample $\tilde{S}$<br>sample $S$ | 1-20 | 21-50 | 51-100 | 101-200 | 201-500 | 501-2000 | 2001+ | Total |
| --- | --- | --- | --- | --- | --- | --- | --- | --- |
| 1-20 | 20 | 0 | 0 | 0 | 0 | 0 | 0 | 20 |
| 21-50 | 0 | 29 | 1 | 0 | 0 | 0 | 0 | 30 |
| 51-100 | 0 | 0 | 39 | 9 | 1 | 0 | 1 | 50 |
| 101-200 | 0 | 1 | 8 | 36 | 14 | 18 | 23 | 100 |
| 201-500 | 0 | 0 | 0 | 26 | 68 | 87 | 119 | 300 |
| 501-2000 | 0 | 0 | 0 | 9 | 101 | 583 | 807 | 1,500 |
| 2001+ | 0 | 0 | 2 | 20 | 116 | 812 | 7,958 | 8,908 |
| Total | 20 | 30 | 50 | 100 | 300 | 1,500 | 8,908 | 10,908 |

For ***A***_*g*_ agreement is better for the top 100 to 500: 99 genes are in the top 100 in both samples, 189 in the top 200; 446 in the top 500 (Table 3).

For ***A***_*g*_ all genes in the top 200 in one sample are in the top 500 in the other. This is not the case for ***S***_*g*_: 42 genes in the top 200 in *S* are not in the top 500 in 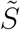; 31 in the top 200 in 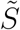 are not in the top 500 in *S*.

For gene *g*, instability is calculated as

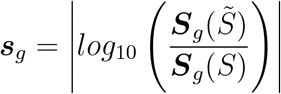

The scores ***S***_*g*_(*S*) and 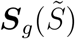 are compared in Figure 6A. In Figure 6B, ***s***_*g*_ is compared to ***S***_*g*_. Figure 6C compares ***s***_*g*_ to ***n***_*g*_ for the 2,000 genes with the largest values of ***S***_*g*_. The Spearman correlation between ***s***_*g*_ and ***n***_*g*_ equals -0.48.

**Figure 6.**
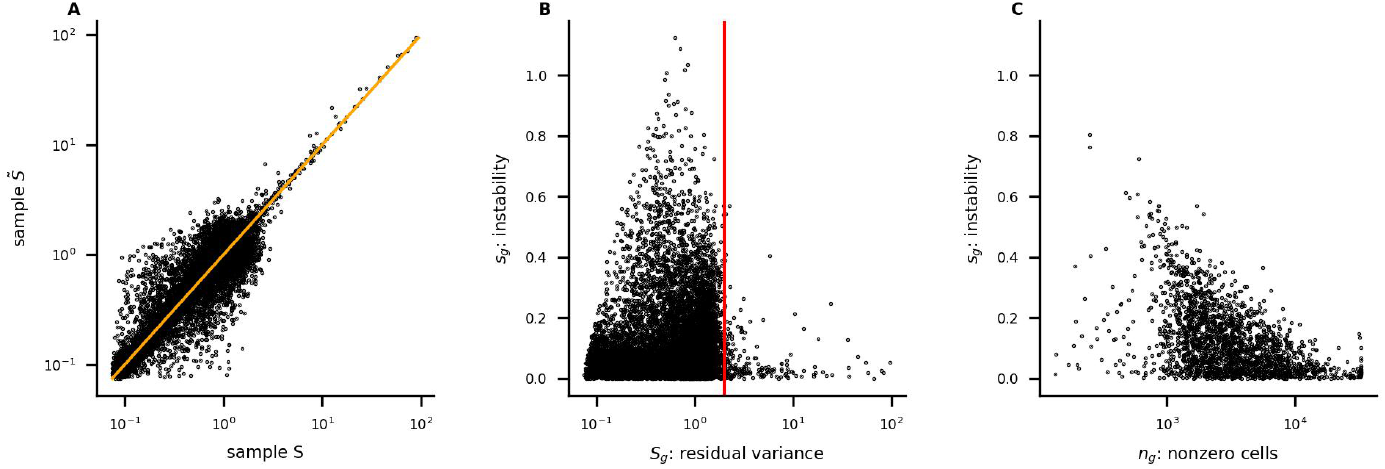
33k PBMC data: (**A**) large values of ***S***_*g*_(*S*) and 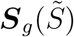 agree closely; (**B**) the red vertical line marks the 200^*th*^ ranked value of ***S***_*g*_ calculated with all cells; (**C**) for the 2,000 genes with the largest values of ***S***_*g*_, instability is larger for genes with fewer nonzero cells (Spearman correlation: -0.48)

As for ***a***_*g*_ and ***l***_*g*_, we compared the distributions of ***a***_*g*_ and ***s***_*g*_ (restricting to genes represented in Figures 2C and 6C) by performing Mann-Whitney and Kolmogorov-Smirnov tests.

The Mann-Whitney one-sided test rejects the null hypothesis that ***s***_*g*_ and ***a***_*g*_ have the same distribution in favor of the alternative that the distribution of ***s***_*g*_ is stochastically greater than that of ***a***_*g*_ (p=0). Similarly, the Kolmogorov-Smirnov one-sided test rejects the null hypothesis in favor of the alternative that the CDF of ***s***_*g*_ is less than that of ***a***_*g*_ (p=0). Repeating the analysis with 19 additional pairs of complementary random samples gave the same results.

### 3.5 Comparing *nru* with *SCTransform* using differential expression analysis

Using Pearson residuals calculated with ***nru*** and ***SCTransform*** and the 10-cluster segmentation from 10x Genomics, Kruskal-Wallis tests were performed for genes with the 2,000 largest values of ***A***_*g*_ and ***S***_*g*_. The methods were compared by:

- plotting ***A***_*g*_ and ***S***_*g*_ vs. the H-statistics for residuals calculated with ***nru*** and ***SC-Transform***, respectively
- selecting the 100 genes ranked highest with each score and comparing the distributions of the corresponding method’s H-statistics, then doing the same for the highest ranked 200, *· · ·* , 1, 000 genes
- tabulating the correlations between each method’s scores and H-statistics

For the 1,133 genes ranked in the top 2,000 by both methods, H-statistics for the two methods agree closely (Figure 7A). Figures 7B and 7C compare ***S***_*g*_ and ***A***_*g*_ to the H-statistics. They suggest that large H-statistics are more likely to be associated with large values of ***A***_*g*_ than ***S***_*g*_. In particular, the vacant region in the lower right corner of Figure 7C shows that no genes have large H-statistics and small values of ***A***_*g*_. This is not the case for ***S***_*g*_ (7B). The gene MALAT1, mentioned in section 3.4, is notated. Its H-statistics are among the largest for both methods.

**Figure 7.**
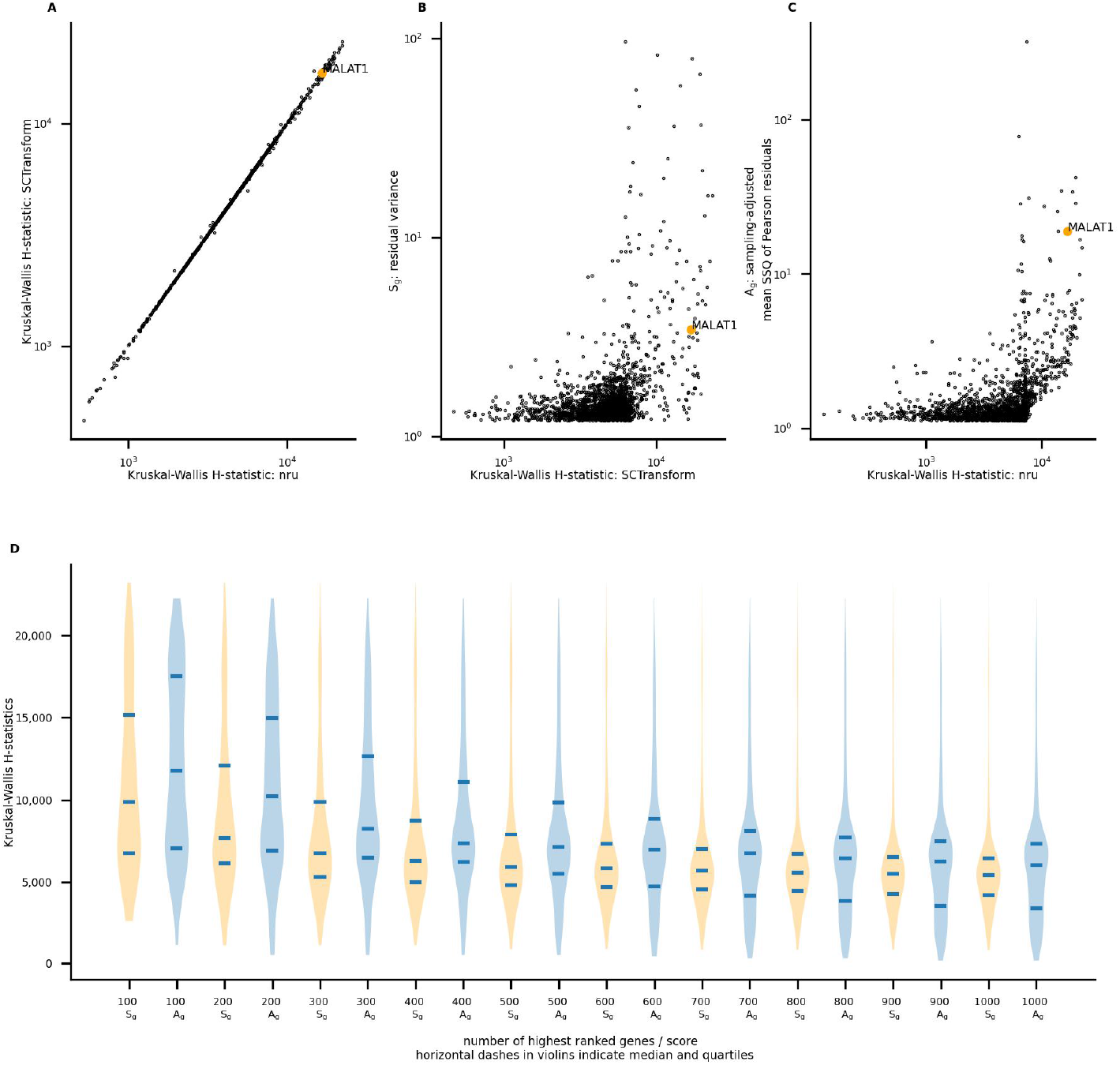
33k PBMC data: (**A**) 1,133 genes are in the highest ranked 2,000 for both methods, the Spearman correlation between their H-statistics equals 1.0; (**B**,**C**) ***A***_*g*_ is large but ***S***_*g*_ is small for several genes with large H-statistics; (**D**) for genes with high scores, H-statistics are generally larger for ***nru***

The violin plots (7D) compare the distributions of H-statistics for residuals calculated with ***nru*** and ***SCTransform*** for the corresponding method’s highest ranked 100, *· · ·*, 1, 000 genes. For all ten pairs of sets of genes, the medians and 75^*th*^ percentiles are larger for ***nru***. For the sets of 200 genes or more, the one-sided Mann-Whitney test rejects the null hypothesis that the distributions of H-statistics for ***nru*** and ***SCTransform*** are equal, in favor of the alternative that the distribution for ***nru*** is stochastically greater (*p <* 0.01). Kolmogorov-Smirnov one-sided tests reject the null hypothesis in favor of the alternative that the CDF of H-statistics for ***nru*** is less than the CDF of the H-statistics for ***SCTransform*** (*p <* 0.003). We note that for the sets of 700 genes or more, one-sided KS tests *also* reject the null hypothesis in favor of the alternative that the CDF of H-statistics for ***nru*** is *greater* than the CDF of the H-statistics for ***SCTransform*** (*p <* 0.003).

Correlations between H-statistics and ***S***_*g*_ or ***A***_*g*_ were calculated for varying numbers of genes with the highest scores. As shown in Table 8, for the sets of 100 to 1,000 genes correlations between ***A***_*g*_ and H-statistics calculated with ***nru*** are larger.

**Table 8.** 33k PBMC data: Spearman correlations between H-statistics and ***S***_*g*_ or ***A***_*g*_ for genes with the largest 50, 100, 200, 500, 1,000, and 2,000 H-statistics.

| | $S_g$ | $A_g$ |
| --- | --- | --- |
| genes |  |  |
| 50 | 0.57 | 0.63 |
| 100 | 0.17 | 0.40 |
| 200 | 0.38 | 0.71 |
| 500 | 0.51 | 0.73 |
| 1000 | 0.39 | 0.51 |
| 2000 | 0.34 | 0.32 |

## 4 Conclusions

We propose normalizing scRNA-seq UMI count data as Pearson residuals of a Poisson model and using the mean sum of squares of the residuals to score genes’ variability. We show that these scores can be unstable if only a few cells are responsible for most of a gene’s total count; such a gene’s data may be unreliable. Variability among scores calculated with random samples of cells identifies many of these genes so that they may be excluded from downstream analysis.

The approach is simple and transparent. It has been observed to identify many of the same highly variable genes as ***scanpy*** and ***SCTransform***. Comparing genes’ scores calculated with UMI count matrices containing half of the cells shows that ***nru*** gives more stable results than the other two methods.

In four of the five data sets that we studied, statistics quantifying differential expression for highly ranked genes are stochastically greater for ***nru*** than ***SCTransform***.

A closed-form expression for the mean sum of squares of Pearson residuals of the Poisson model provides insights: only the cells in which a gene’s counts are nonzero contribute; each cell’s contribution depends on its sequencing depth, the gene’s count in the cell, and the UMI matrix’s total count and number of cells.

## A A closed-form expression for the mean sum of squares of Pearson residuals of a Poisson model

We use the notations

*X*_*gc*_ = the UMI count for gene *g* in cell *c*

*G*_*g*_ = ∑_*c*_ *X*_*gc*_ the total count for gene *g*

*π*_*c*|*g*_ = *X*_*gc*_*/G*_*g*_ the fraction of the total count of gene *g* that is in cell *c*

*D*_*c*_ =∑ _*g*_ *X*_*gc*_ the total count for cell *c* – its sequencing depth

*T* = ∑_*gc*_ *X*_*gc*_ the total of all counts in the matrix

*C* = the total number of cells in the matrix

*D* = *T/C* the mean sequencing depth of the matrix

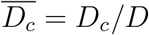 the normalized sequencing depth of cell *c*

The Pearson residuals are

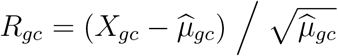

with the maximum likelihood estimate

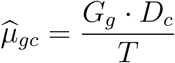

Squaring gives

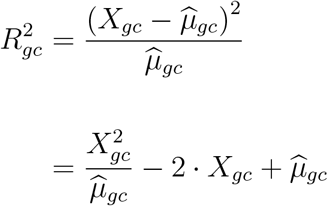

Summing over all cells *c* gives ***P*** _*g*_, the sum of squares of Pearson residuals for gene *g*:

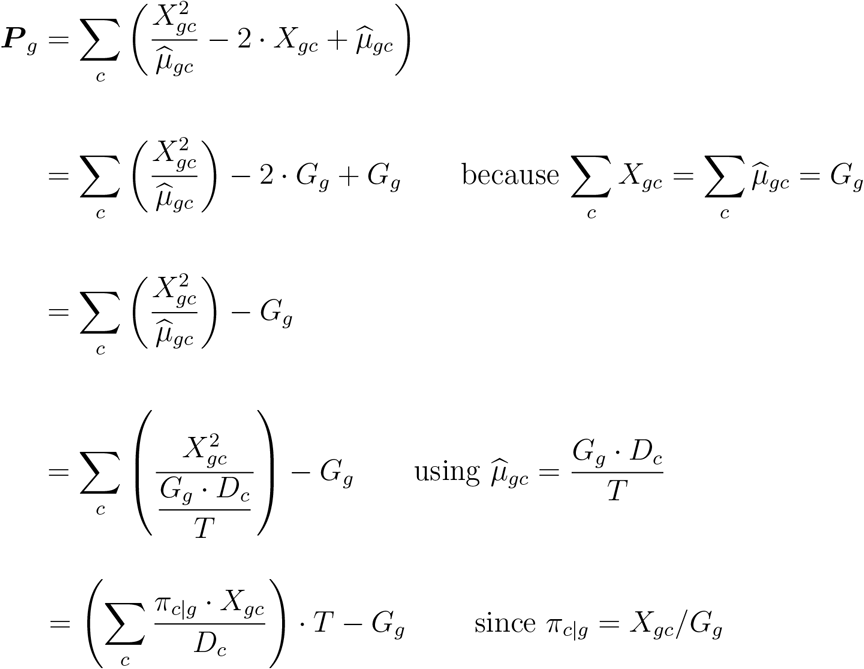

Dividing by the number of cells in the UMI matrix, *C*, gives the mean sum of squares of Pearson residuals for gene *g*:

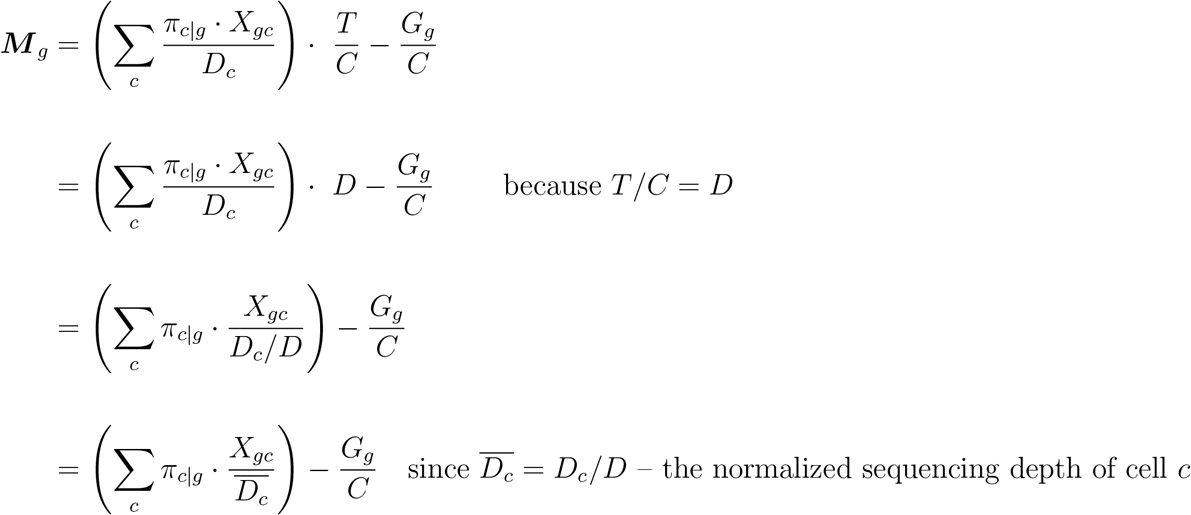

Observations:

- The summation includes *only* cells on which gene *g* has nonzero counts.
- The mean value of the denominators 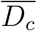 *over all cells in the UMI count matrix* is 1.
- The subtracted term 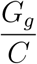 is negligible for lowly expressed genes.
- Since ∑*π*_*c*|*g*_ = 1, the summation is the convex combination of the ratios 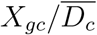. This provides intuition for the fact that many values of ***M*** _*g*_ are near 1. They are convex combinations of terms that are often close to 1: most nonzero counts *X*_*gc*_ equal 1 and the mean of the 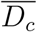 equals 1.

