## Supplement 1 for "Normalization and gene selection for single-cell RNA-seq UMI data using sampling-adjusted sums of squares of Pearson residuals with a Poisson model"

We include figures and tables corresponding to those in the Results and discussion section of the paper for four additional data sets: retinal, lupus, 10k brain, and 10k heart.

Our programs, in Jupyter notebooks containing these results, are available at https://github.com/victorkleb/scRNA-seq_Pr

For the heart data set, we discuss the three genes for which ***nru***'s results calculated with complementary random samples of half of the cells are unstable. For these genes, counts are concentrated on a small number of cells, suggesting that this may identify a quality control issue.

### retinal data

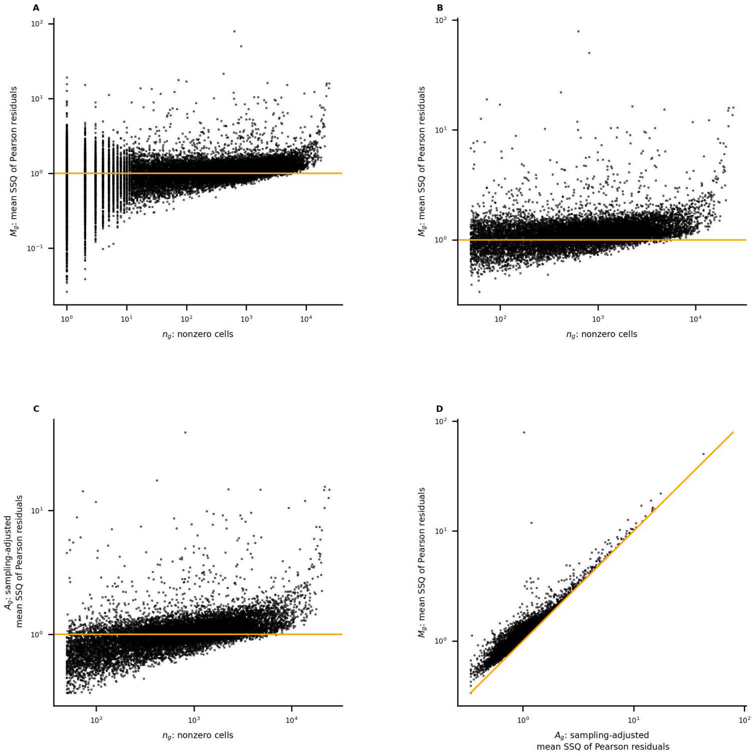


**Figure R-1** retinal data: (**A**) the mean SSQ of Pearson residuals $\boldsymbol{M}_g$ vs. the number of nonzero cells $\boldsymbol{n}_g$ for all genes; (**B**) restricting to genes with at least 50 nonzero cells; (**C**) $\boldsymbol{A}_g$ vs. $\boldsymbol{n}_g$; (**D**) the effect of sampling-adjustment: $\boldsymbol{M}_g$ vs. $\boldsymbol{A}_g$

| $\boldsymbol{M}_g$ <br> $\boldsymbol{n}_g$ | $\leq 1$ | $1 < -2$ | $> 2$ | Total |
|---|---|---|---|---|
| 1-10 | 3,725 | 1,712 | 507 | 5,944 |
| 11-50 | 1,693 | 1,077 | 67 | 2,837 |
| 51-100 | 665 | 612 | 25 | 1,302 |
| 101-1,000 | 2,090 | 4,368 | 109 | 6,567 |
| 1,001-10,000 | 453 | 4,908 | 147 | 5,508 |
| 10,001+ | 0 | 62 | 72 | 134 |
| Total | 8,626 | 12,739 | 927 | 22,292 |

**Table R-1** retinal data: relation between $\boldsymbol{n}_g$, the number of nonzero cells, and $\boldsymbol{M}_g$, the mean SSQ of Pearson residuals

| sample $\tilde{S}$ <br> sample $S$ | 1-20 | 21-50 | 51-100 | 101-200 | 201-500 | 501-2000 | 2001+ | Total |
|---|---|---|---|---|---|---|---|---|
| 1-20 | 18 | 2 | 0 | 0 | 0 | 0 | 0 | 20 |
| 21-50 | 2 | 25 | 3 | 0 | 0 | 0 | 0 | 30 |
| 51-100 | 0 | 3 | 42 | 5 | 0 | 0 | 0 | 50 |
| 101-200 | 0 | 0 | 5 | 81 | 12 | 1 | 1 | 100 |
| 201-500 | 0 | 0 | 0 | 14 | 224 | 61 | 1 | 300 |
| 501-2000 | 0 | 0 | 0 | 0 | 62 | 1,109 | 329 | 1,500 |
| 2001+ | 0 | 0 | 0 | 0 | 2 | 329 | 9,751 | 10,082 |
| Total | 20 | 30 | 50 | 100 | 300 | 1,500 | 10,082 | 12,082 |

**Table R-2** retinal data: comparing ranks of $\boldsymbol{A}_g$ calculated with $S$ and $\tilde{S}$

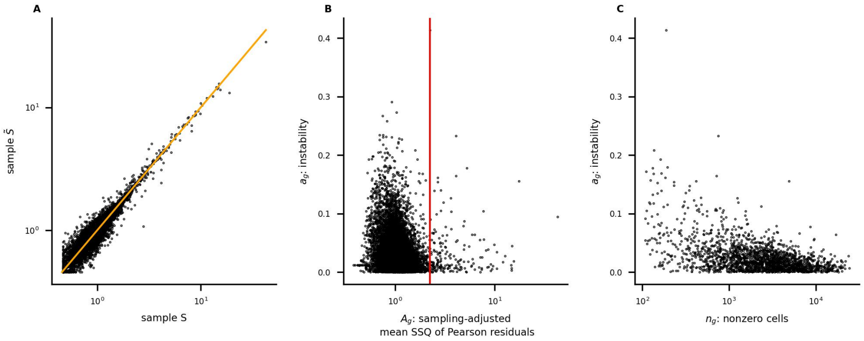


**Figure R-2** retinal data: (**A**) large values of $\boldsymbol{A}_g(S)$ and $\boldsymbol{A}_g(\tilde{S})$ agree closely; (**B**) the red vertical line marks the $200^{th}$ ranked value of $\boldsymbol{A}_g$ calculated with all cells; (**C**) for genes with the 2,000 largest values of $\boldsymbol{A}_g$, for which both $\boldsymbol{A}_g(S)$ and $\boldsymbol{A}_g(\tilde{S})$ were also calculated, instability is larger for genes with fewer nonzero cells

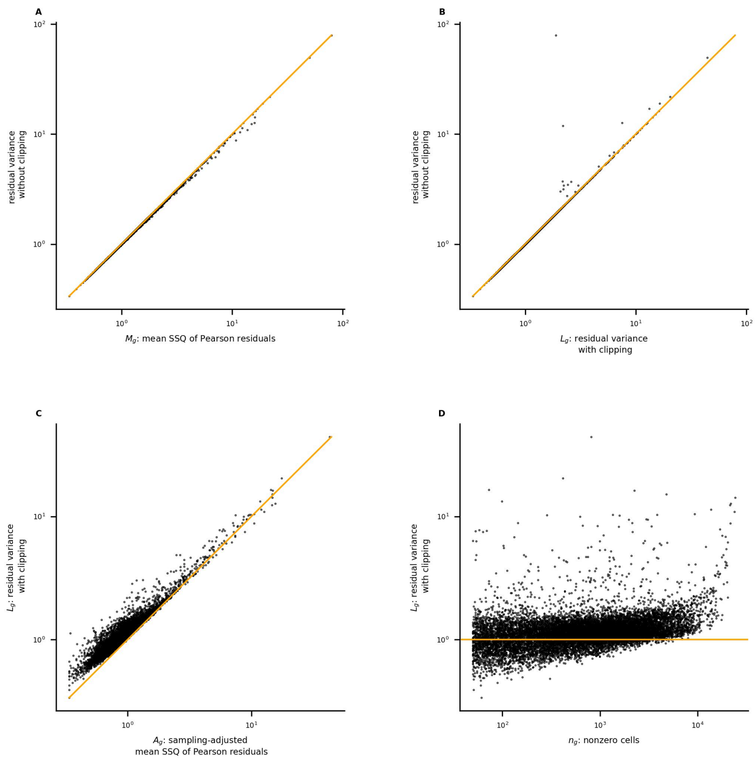


**Figure R-3** retinal data: (**A**) scores calculated by ***scanpy*** without clipping agree closely with $\boldsymbol{M}_g$; (**B**) clipping affects scores for very few genes; (**C**) $\boldsymbol{L}_g$ and $\boldsymbol{A}_g$ are large for many of the same genes; (**D**) ***scanpy*** score $\boldsymbol{L}_g$ vs. the number of nonzero cells

| $\boldsymbol{L}_g$ rank<br>$\boldsymbol{A}_g$ rank | 1-20 | 21-50 | 51-100 | 101-200 | 201-500 | 501+ | Total |
|---|---|---|---|---|---|---|---|
| 1-20 | 18 | 2 | 0 | 0 | 0 | 0 | 20 |
| 21-50 | 2 | 23 | 5 | 0 | 0 | 0 | 30 |
| 51-100 | 0 | 5 | 36 | 9 | 0 | 0 | 50 |
| 101-200 | 0 | 0 | 9 | 68 | 23 | 0 | 100 |
| 201-500 | 0 | 0 | 0 | 17 | 213 | 70 | 300 |
| 501+ | 0 | 0 | 0 | 6 | 64 | 12,982 | 13,052 |
| Total | 20 | 30 | 50 | 100 | 300 | 13,052 | 13,552 |

**Table R-3** retinal data: comparing ranks of $\boldsymbol{L}_g$ with $\boldsymbol{A}_g$

| sample $\tilde{S}$<br>sample $S$ | 1-20 | 21-50 | 51-100 | 101-200 | 201-500 | 501-2000 | 2001+ | Total |
|---|---|---|---|---|---|---|---|---|
| 1-20 | 19 | 1 | 0 | 0 | 0 | 0 | 0 | 20 |
| 21-50 | 1 | 27 | 2 | 0 | 0 | 0 | 0 | 30 |
| 51-100 | 0 | 2 | 40 | 7 | 1 | 0 | 0 | 50 |
| 101-200 | 0 | 0 | 6 | 78 | 14 | 2 | 0 | 100 |
| 201-500 | 0 | 0 | 2 | 14 | 209 | 60 | 15 | 300 |
| 501-2000 | 0 | 0 | 0 | 1 | 70 | 1,000 | 429 | 1,500 |
| 2001+ | 0 | 0 | 0 | 0 | 6 | 438 | 9,638 | 10,082 |
| Total | 20 | 30 | 50 | 100 | 300 | 1,500 | 10,082 | 12,082 |

**Table R-4** retinal data: comparing ranks of $\boldsymbol{L}_g$ calculated with $S$ and $\tilde{S}$

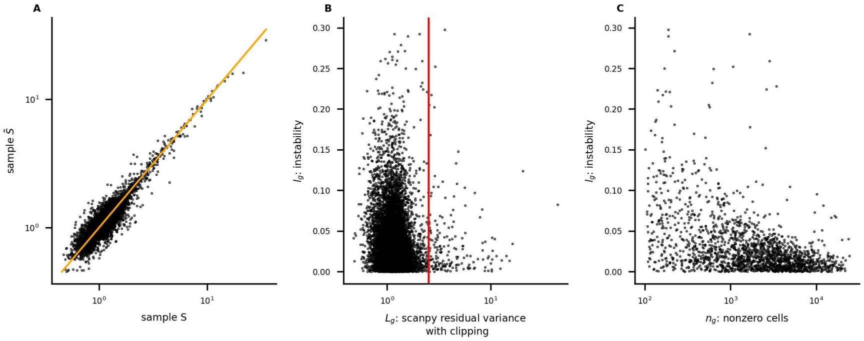


**Figure R-4** retinal data: (**A**) large values of $\boldsymbol{L}_g(S)$ and $\boldsymbol{L}_g(\tilde{S})$ agree closely; (**B**) the red vertical line marks the $200^{th}$ ranked value of $\boldsymbol{L}_g$ calculated with all cells; (**C**) for genes with the 2,000 largest values of $\boldsymbol{L}_g$, for which both $\boldsymbol{L}_g(S)$ and $\boldsymbol{L}_g(\tilde{S})$ were also calculated, instability is larger for genes with fewer nonzero cells

The distributions of $\boldsymbol{a}_g$ and $\boldsymbol{l}_g$ were compared by performing Mann-Whitney and Kolmogorov-Smirnov tests (restricting to genes represented in Figures R-2C and R-4C).

The Mann-Whitney one-sided test rejects the null hypothesis that $\boldsymbol{l}_g$ and $\boldsymbol{a}_g$ have the same distribution in favor of the alternative that the distribution of $\boldsymbol{l}_g$ is stochastically greater than that of $\boldsymbol{a}_g$ (p=0.008). Similarly, the Kolmogorov-Smirnov one-sided test rejects the null hypothesis in favor of the alternative that the CDF of $\boldsymbol{l}_g$ is less than that of $\boldsymbol{a}_g$ (p=0.0008).

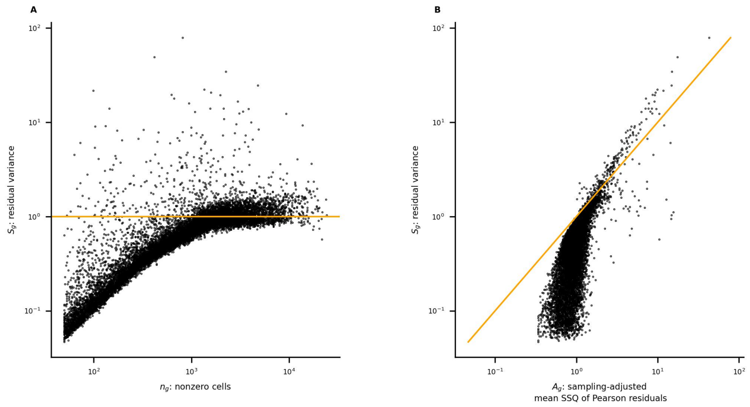


**Figure R-5** retinal data: (**A**) ***SCTransform*** assigns very low scores to most genes with few nonzero cells; (**B**) $\boldsymbol{S}_g$ and $\boldsymbol{A}_g$ are large for many of the same genes

| $\boldsymbol{S}_g$ rank<br>$\boldsymbol{A}_g$ rank | 1-20 | 21-50 | 51-100 | 101-200 | 201-500 | 501+ | Total |
|---|---|---|---|---|---|---|---|
| 1-20 | 12 | 2 | 1 | 0 | 1 | 4 | 20 |
| 21-50 | 8 | 13 | 1 | 2 | 2 | 4 | 30 |
| 51-100 | 0 | 15 | 18 | 4 | 6 | 7 | 50 |
| 101-200 | 0 | 0 | 30 | 42 | 20 | 8 | 100 |
| 201-500 | 0 | 0 | 0 | 45 | 182 | 73 | 300 |
| 501+ | 0 | 0 | 0 | 7 | 89 | 12,956 | 13,052 |
| Total | 20 | 30 | 50 | 100 | 300 | 13,052 | 13,552 |

**Table R-5** retinal data: comparing ranks of $\boldsymbol{S}_g$ with $\boldsymbol{A}_g$

| sample $\tilde{S}$<br>sample $S$ | 1-20 | 21-50 | 51-100 | 101-200 | 201-500 | 501-2000 | 2001+ | Total |
|---|---|---|---|---|---|---|---|---|
| 1-20 | 20 | 0 | 0 | 0 | 0 | 0 | 0 | 20 |
| 21-50 | 0 | 28 | 1 | 0 | 1 | 0 | 0 | 30 |
| 51-100 | 0 | 2 | 40 | 8 | 0 | 0 | 0 | 50 |
| 101-200 | 0 | 0 | 9 | 73 | 14 | 2 | 2 | 100 |
| 201-500 | 0 | 0 | 0 | 15 | 208 | 72 | 5 | 300 |
| 501-2000 | 0 | 0 | 0 | 3 | 73 | 1,203 | 221 | 1,500 |
| 2001+ | 0 | 0 | 0 | 1 | 4 | 223 | 9,854 | 10,082 |
| Total | 20 | 30 | 50 | 100 | 300 | 1,500 | 10,082 | 12,082 |

**Table R-6** retinal data: comparing ranks of $\boldsymbol{S}_g$ calculated with $S$ and $\tilde{S}$

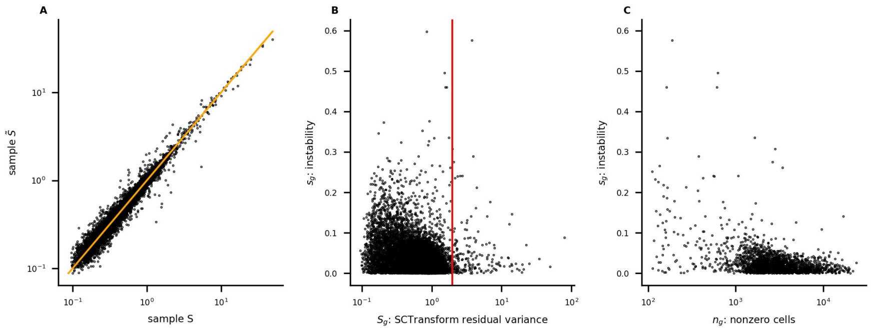


**Figure R-6** retinal data: (**A**) large values of $\boldsymbol{S}_g(S)$ and $\boldsymbol{S}_g(\tilde{S})$ agree closely; (**B**) the red vertical line marks the $200^{th}$ ranked value of $\boldsymbol{S}_g$ calculated with all cells; (**C**) for genes with the 2,000 largest values of $\boldsymbol{S}_g$, for which both $\boldsymbol{S}_g(S)$ and $\boldsymbol{S}_g(\tilde{S})$ were also calculated, instability is larger for genes with fewer nonzero cells

The distributions of $\boldsymbol{a}_g$ and $\boldsymbol{s}_g$ were compared by performing Mann-Whitney and Kolmogorov-Smirnov tests (restricting to genes represented in Figures R-2C and R-6C).

The Mann-Whitney one-sided test rejects the null hypothesis that $\boldsymbol{s}_g$ and $\boldsymbol{a}_g$ have the same distribution in favor of the alternative that the distribution of $\boldsymbol{s}_g$ is stochastically greater than that of $\boldsymbol{a}_g$ (p=0.02). Similarly, the Kolmogorov-Smirnov one-sided test rejects the null hypothesis in favor of the alternative that the CDF of $\boldsymbol{s}_g$ is less than that of $\boldsymbol{a}_g$ (p=0.05).

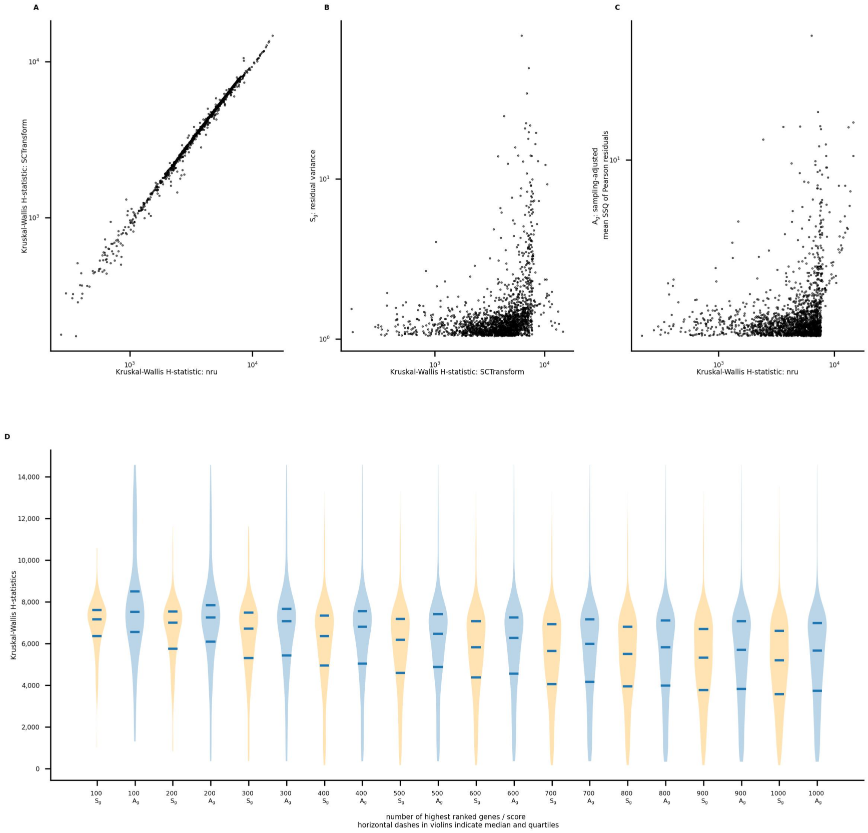


**Figure R-7** retinal data: Kruskal-Wallis H-statistics were calculated using a 39-cluster segmentation; (**A**) for genes ranked in the top 2,000 by both methods, H-statistics are strongly correlated; (**B,C**) $\boldsymbol{A}_g$ is large but $\boldsymbol{S}_g$ is small for several genes with large H-statistics; (**D**) for genes with high scores, H-statistics are generally larger for ***nru***

Using Pearson residuals calculated with ***nru*** and ***SCTransform***, Kruskal-Wallis tests were performed for genes with the 2,000 largest values of $\boldsymbol{A}_g$ and $\boldsymbol{S}_g$.

The violin plots (R-7D) compare the distributions of H-statistics for residuals calculated with ***nru*** and ***SCTransform*** for the corresponding method's highest ranked $100, \cdots, 1,000$ genes.

For each pair of sets of genes, the one-sided Mann-Whitney test rejects the null hypothesis that the distributions of H-statistics for ***nru*** and ***SCTransform*** are equal, in favor of the alternative that the distribution for ***nru*** is stochastically greater ($p < 0.02$). Kolmogorov-Smirnov one-sided tests reject the null hypothesis in favor of the alternative that the CDF of H-statistics for ***nru*** is less than the CDF of the H-statistics for ***SCTransform*** ($p < 0.02$).

| genes | $\boldsymbol{S}_g$ | $\boldsymbol{A}_g$ |
|---|---|---|
| 50 | -0.49 | 0.52 |
| 100 | -0.09 | 0.42 |
| 200 | 0.08 | 0.61 |
| 500 | 0.38 | 0.42 |
| 1000 | 0.48 | 0.31 |
| 2000 | 0.42 | 0.21 |

**Table R-7** retinal data: Spearman correlations between H-statistics and $\boldsymbol{S}_g$ or $\boldsymbol{A}_g$ for genes with the largest 50, 100, 200, 500, 1,000, and 2,000 H-statistics

For the sets of genes with the largest 50, 100, 200, 500, 1,000, and 2,000 H-statistics, Spearman correlations between H-statistics and $\boldsymbol{S}_g$ or $\boldsymbol{A}_g$ were calculated. They are summarized in Table R-7. For the sets of 50, 100, and 200 genes, correlations between $\boldsymbol{A}_g$ and H-statistics calculated with $\boldsymbol{nru}$ are larger.

### lupus data

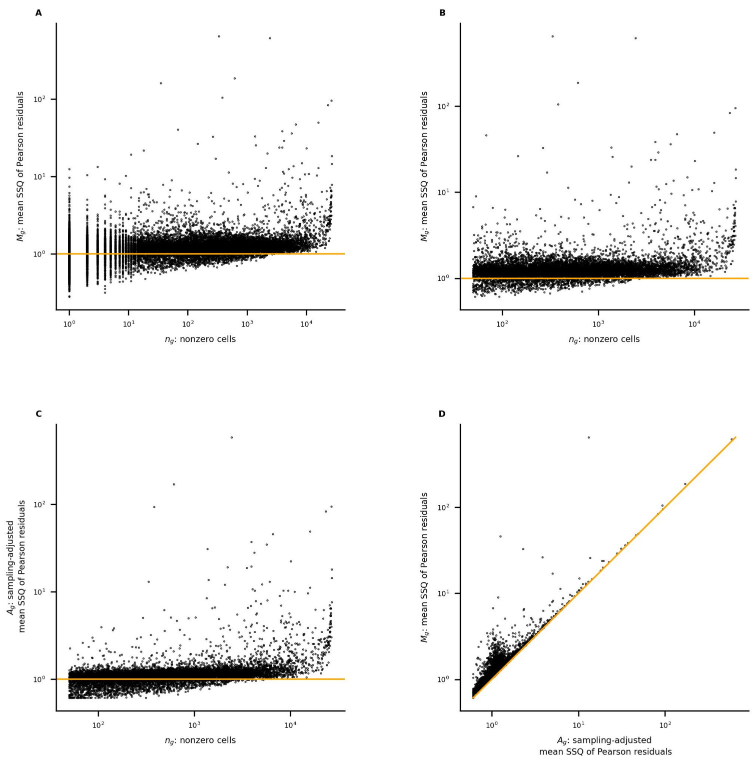


**Figure L-1** lupus data: (**A**) the mean SSQ of Pearson residuals $\boldsymbol{M}_g$ vs. the number of nonzero cells $\boldsymbol{n}_g$ for all genes; (**B**) restricting to genes with at least 50 nonzero cells; (**C**) $\boldsymbol{A}_g$ vs. $\boldsymbol{n}_g$; (**D**) the effect of sampling-adjustment: $\boldsymbol{M}_g$ vs. $\boldsymbol{A}_g$

| $\boldsymbol{M}_g$ | $\leq 1$ | $1 < -2$ | $> 2$ | Total |
|---|---|---|---|---|
| $\boldsymbol{n}_g$ | | | | |
| 1-10 | 2,027 | 3,009 | 232 | 5,268 |
| 11-50 | 563 | 1,995 | 89 | 2,647 |
| 51-100 | 212 | 1,085 | 55 | 1,352 |
| 101-1,000 | 640 | 5,233 | 119 | 5,992 |
| 1,001-10,000 | 75 | 2,788 | 147 | 3,010 |
| 10,001+ | 0 | 115 | 117 | 232 |
| Total | 3,517 | 14,225 | 759 | 18,501 |

**Table L-1** lupus data: relation between $\boldsymbol{n}_g$, the number of nonzero cells, and $\boldsymbol{M}_g$, the mean SSQ of Pearson residuals

| sample $\tilde{S}$<br>sample $S$ | 1-20 | 21-50 | 51-100 | 101-200 | 201-500 | 501-2000 | 2001+ | Total |
|---|---|---|---|---|---|---|---|---|
| 1-20 | 18 | 2 | 0 | 0 | 0 | 0 | 0 | 20 |
| 21-50 | 2 | 25 | 2 | 1 | 0 | 0 | 0 | 30 |
| 51-100 | 0 | 1 | 45 | 3 | 1 | 0 | 0 | 50 |
| 101-200 | 0 | 2 | 3 | 87 | 8 | 0 | 0 | 100 |
| 201-500 | 0 | 0 | 0 | 9 | 250 | 41 | 0 | 300 |
| 501-2000 | 0 | 0 | 0 | 0 | 39 | 986 | 475 | 1,500 |
| 2001+ | 0 | 0 | 0 | 0 | 2 | 473 | 6,616 | 7,091 |
| Total | 20 | 30 | 50 | 100 | 300 | 1,500 | 7,091 | 9,091 |

**Table L-2** lupus data: comparing ranks of $\boldsymbol{A}_g$ calculated with $S$ and $\tilde{S}$

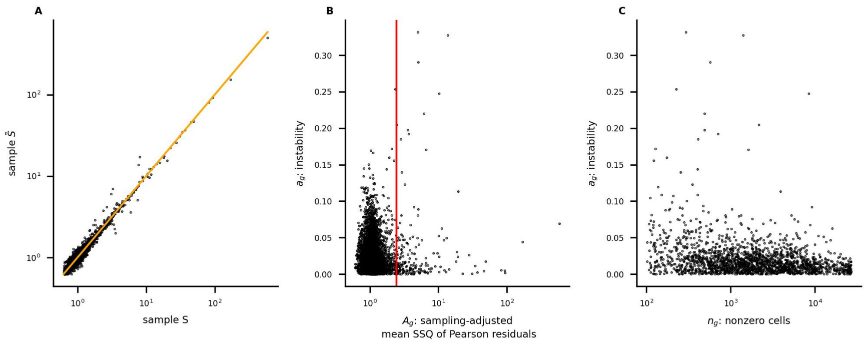


**Figure L-2** lupus data: (**A**) large values of $\boldsymbol{A}_g(S)$ and $\boldsymbol{A}_g(\tilde{S})$ agree closely; (**B**) the red vertical line marks the $200^{th}$ ranked value of $\boldsymbol{A}_g$ calculated with all cells; (**C**) for genes with the 2,000 largest values of $\boldsymbol{A}_g$, for which both $\boldsymbol{A}_g(S)$ and $\boldsymbol{A}_g(\tilde{S})$ were also calculated, instability is larger for genes with fewer nonzero cells

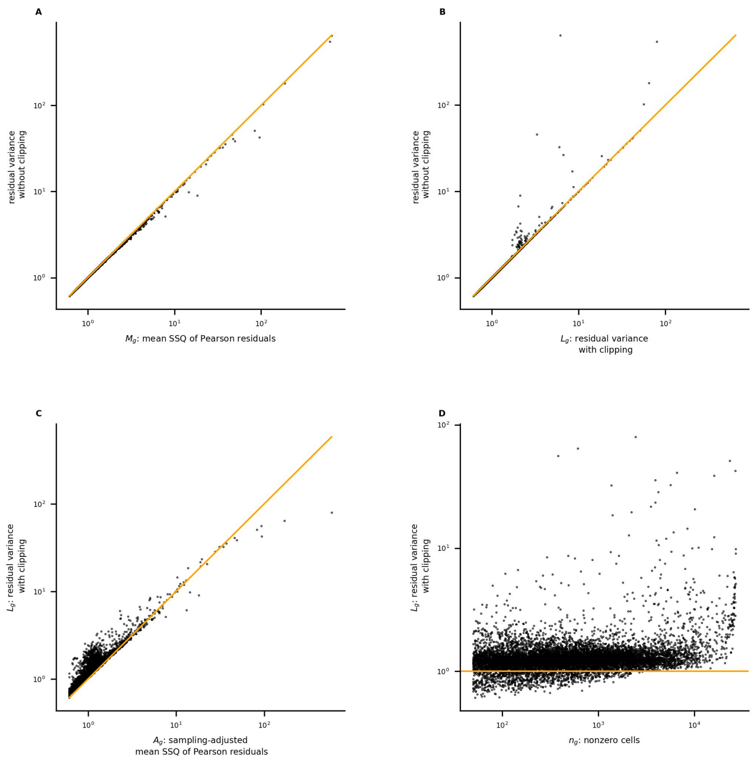


**Figure L-3** lupus data: (**A**) scores calculated by ***scanpy*** without clipping agree closely with $\boldsymbol{M}_g$; (**B**) clipping affects scores for very few genes; (**C**) $\boldsymbol{L}_g$ and $\boldsymbol{A}_g$ are large for many of the same genes; (**D**) ***scanpy*** score $\boldsymbol{L}_g$ vs. the number of nonzero cells

| $\boldsymbol{L}_g$ rank<br>$\boldsymbol{A}_g$ rank | 1-20 | 21-50 | 51-100 | 101-200 | 201-500 | 501+ | Total |
|---|---|---|---|---|---|---|---|
| 1-20 | 17 | 3 | 0 | 0 | 0 | 0 | 20 |
| 21-50 | 3 | 19 | 8 | 0 | 0 | 0 | 30 |
| 51-100 | 0 | 7 | 30 | 13 | 0 | 0 | 50 |
| 101-200 | 0 | 0 | 9 | 64 | 27 | 0 | 100 |
| 201-500 | 0 | 1 | 3 | 17 | 139 | 140 | 300 |
| 501+ | 0 | 0 | 0 | 6 | 134 | 9,979 | 10,119 |
| Total | 20 | 30 | 50 | 100 | 300 | 10,119 | 10,619 |

**Table L-3** lupus data: comparing ranks of $\boldsymbol{L}_g$ with $\boldsymbol{A}_g$

| sample $\tilde{S}$ <br> sample $S$ | 1-20 | 21-50 | 51-100 | 101-200 | 201-500 | 501-2000 | 2001+ | Total |
|---|---|---|---|---|---|---|---|---|
| 1-20 | 18 | 2 | 0 | 0 | 0 | 0 | 0 | 20 |
| 21-50 | 1 | 25 | 2 | 2 | 0 | 0 | 0 | 30 |
| 51-100 | 1 | 3 | 41 | 5 | 0 | 0 | 0 | 50 |
| 101-200 | 0 | 0 | 7 | 74 | 18 | 0 | 1 | 100 |
| 201-500 | 0 | 0 | 0 | 13 | 120 | 53 | 114 | 300 |
| 501-2000 | 0 | 0 | 0 | 2 | 52 | 631 | 815 | 1,500 |
| 2001+ | 0 | 0 | 0 | 4 | 110 | 816 | 6,161 | 7,091 |
| Total | 20 | 30 | 50 | 100 | 300 | 1,500 | 7,091 | 9,091 |

**Table L-4** lupus data: comparing ranks of $\boldsymbol{L}_g$ calculated with $S$ and $\tilde{S}$

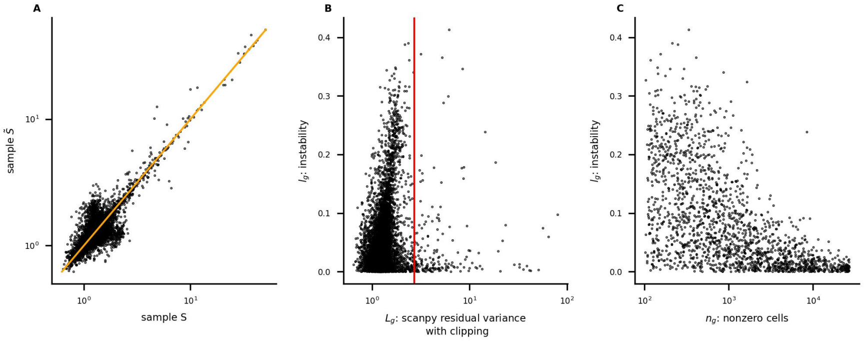


**Figure L-4** lupus data: (**A**) most large values of $\boldsymbol{L}_g(S)$ and $\boldsymbol{L}_g(\tilde{S})$ agree closely; (**B**) the red vertical line marks the $200^{th}$ ranked value of $\boldsymbol{L}_g$ calculated with all cells; (**C**) for genes with the 2,000 largest values of $\boldsymbol{L}_g$, for which both $\boldsymbol{L}_g(S)$ and $\boldsymbol{L}_g(\tilde{S})$ were also calculated, instability is larger for genes with fewer nonzero cells

The distributions of $\boldsymbol{a}_g$ and $\boldsymbol{l}_g$ were compared by performing Mann-Whitney and Kolmogorov-Smirnov tests (restricting to genes represented in Figures L-2C and L-4C).

The Mann-Whitney one-sided test rejects the null hypothesis that $\boldsymbol{l}_g$ and $\boldsymbol{a}_g$ have the same distribution in favor of the alternative that the distribution of $\boldsymbol{l}_g$ is stochastically greater than that of $\boldsymbol{a}_g$ (p=0). Similarly, the Kolmogorov-Smirnov one-sided test rejects the null hypothesis in favor of the alternative that the CDF of $\boldsymbol{l}_g$ is less than that of $\boldsymbol{a}_g$ (p=0).

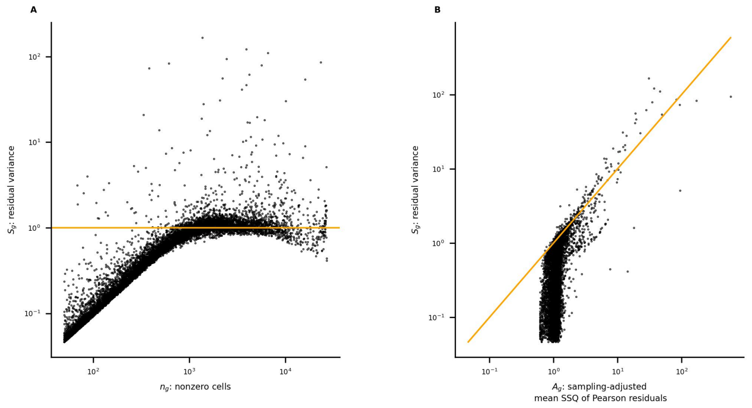


**Figure L-5** lupus data: (**A**) ***SCTransform*** assigns very low scores to most genes with few nonzero cells; (**B**) $\boldsymbol{S}_g$ and $\boldsymbol{A}_g$ are large for many of the same genes

| $\boldsymbol{S}_g$ rank<br>$\boldsymbol{A}_g$ rank | 1-20 | 21-50 | 51-100 | 101-200 | 201-500 | 501+ | Total |
|---|---|---|---|---|---|---|---|
| 1-20 | 17 | 0 | 1 | 0 | 1 | 1 | 20 |
| 21-50 | 3 | 18 | 2 | 4 | 2 | 1 | 30 |
| 51-100 | 0 | 10 | 20 | 7 | 5 | 8 | 50 |
| 101-200 | 0 | 1 | 23 | 26 | 17 | 33 | 100 |
| 201-500 | 0 | 1 | 3 | 61 | 111 | 124 | 300 |
| 501+ | 0 | 0 | 1 | 2 | 164 | 9,952 | 10,119 |
| Total | 20 | 30 | 50 | 100 | 300 | 10,119 | 10,619 |

**Table L-5** lupus data: comparing ranks of $\boldsymbol{S}_g$ with $\boldsymbol{A}_g$

| sample $\tilde{S}$<br>sample $S$ | 1-20 | 21-50 | 51-100 | 101-200 | 201-500 | 501-2000 | 2001+ | Total |
|---|---|---|---|---|---|---|---|---|
| 1-20 | 19 | 1 | 0 | 0 | 0 | 0 | 0 | 20 |
| 21-50 | 1 | 26 | 3 | 0 | 0 | 0 | 0 | 30 |
| 51-100 | 0 | 3 | 39 | 6 | 1 | 0 | 1 | 50 |
| 101-200 | 0 | 0 | 7 | 70 | 20 | 3 | 0 | 100 |
| 201-500 | 0 | 0 | 1 | 17 | 151 | 108 | 23 | 300 |
| 501-2000 | 0 | 0 | 0 | 4 | 106 | 1,044 | 346 | 1,500 |
| 2001+ | 0 | 0 | 0 | 3 | 22 | 345 | 6,721 | 7,091 |
| Total | 20 | 30 | 50 | 100 | 300 | 1,500 | 7,091 | 9,091 |

**Table L-6** lupus data: comparing ranks of $\boldsymbol{S}_g$ calculated with $S$ and $\tilde{S}$

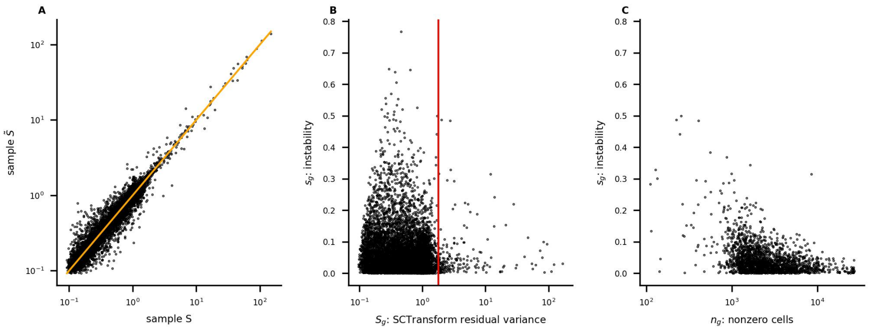


**Figure L-6** lupus data: (**A**) large values of $\boldsymbol{S}_g(S)$ and $\boldsymbol{S}_g(\tilde{S})$ agree closely; (**B**) the red vertical line marks the $200^{th}$ ranked value of $\boldsymbol{S}_g$ calculated with all cells; (**C**) for genes with the 2,000 largest values of $\boldsymbol{S}_g$, for which both $\boldsymbol{S}_g(S)$ and $\boldsymbol{S}_g(\tilde{S})$ were also calculated, instability is larger for genes with fewer nonzero cells

The distributions of $\boldsymbol{a}_g$ and $\boldsymbol{s}_g$ were compared by performing Mann-Whitney and Kolmogorov-Smirnov tests (restricting to genes represented in Figures L-2C and L-6C).

The Mann-Whitney one-sided test rejects the null hypothesis that $\boldsymbol{s}_g$ and $\boldsymbol{a}_g$ have the same distribution in favor of the alternative that the distribution of $\boldsymbol{s}_g$ is stochastically greater than that of $\boldsymbol{a}_g$ (p=0). Similarly, the Kolmogorov-Smirnov one-sided test rejects the null hypothesis in favor of the alternative that the CDF of $\boldsymbol{s}_g$ is less than that of $\boldsymbol{a}_g$ (p=0).

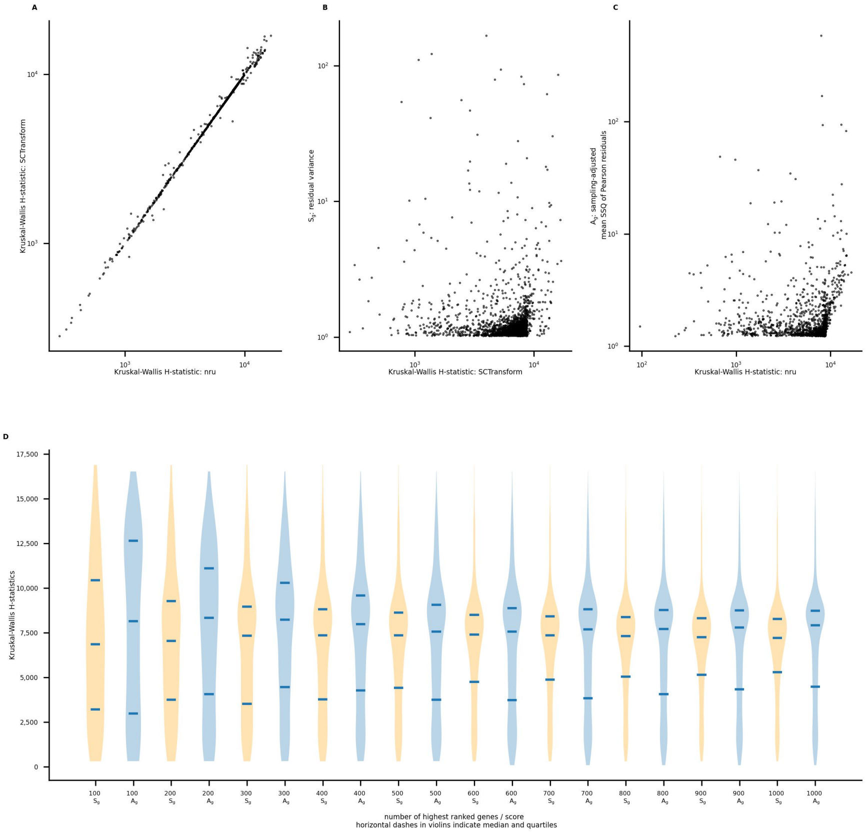


**Figure L-7** lupus data: Kruskal-Wallis H-statistics were calculated using an 8-cluster segmentation; (**A**) for genes ranked in the top 2,000 by both methods, H-statistics are strongly correlated; (**B**) $\boldsymbol{S_g}$ is small for several genes with large H-statistics; (**D**) for genes with high scores, H-statistics are generally larger for ***nru***

The violin plots (L-7D) compare the distributions of H-statistics for residuals calculated with ***nru*** and ***SCTransform*** for the corresponding method's highest ranked $100, \cdots, 1,000$ genes.

For the sets of 300, 400, 800, 900, and 1000 genes, the one-sided Mann-Whitney test rejects the null hypothesis that the distributions of H-statistics for ***nru*** and ***SCTransform*** are equal, in favor of the alternative that the distribution for ***nru*** is stochastically greater ($p < 0.02$). For the sets of 300 genes or more, Kolmogorov-Smirnov one-sided tests reject the null hypothesis in favor of the alternative that the CDF of H-statistics for ***nru*** is less than the CDF of the H-statistics for ***SCTransform*** ($p < 0.003$). We note that for the sets of 600 genes or more, one-sided KS tests *also* reject the null hypothesis in favor of the alternative that the CDF of H-statistics for ***nru*** is *greater* than the CDF of the H-statistics for ***SCTransform*** ($p < 0.05$).

| genes | $\boldsymbol{S}_g$ | $\boldsymbol{A}_g$ |
|---|---|---|
| 50 | 0.51 | 0.38 |
| 100 | 0.34 | 0.70 |
| 200 | 0.50 | 0.86 |
| 500 | 0.46 | 0.75 |
| 1000 | 0.37 | 0.45 |
| 2000 | 0.16 | -0.05 |

**Table L-7** lupus data: Spearman correlations between H-statistics and $\boldsymbol{S}_g$ or $\boldsymbol{A}_g$ for genes with the largest 50, 100, 200, 500, 1,000, and 2,000 H-statistics

Correlations between H-statistics and $\boldsymbol{S}_g$ or $\boldsymbol{A}_g$ were calculated for varying numbers of genes with the highest scores. For the sets of 100, 200, and 500 genes, correlations between $\boldsymbol{A}_g$ and H-statistics calculated with $\boldsymbol{nru}$ are larger.

#### 10k brain data

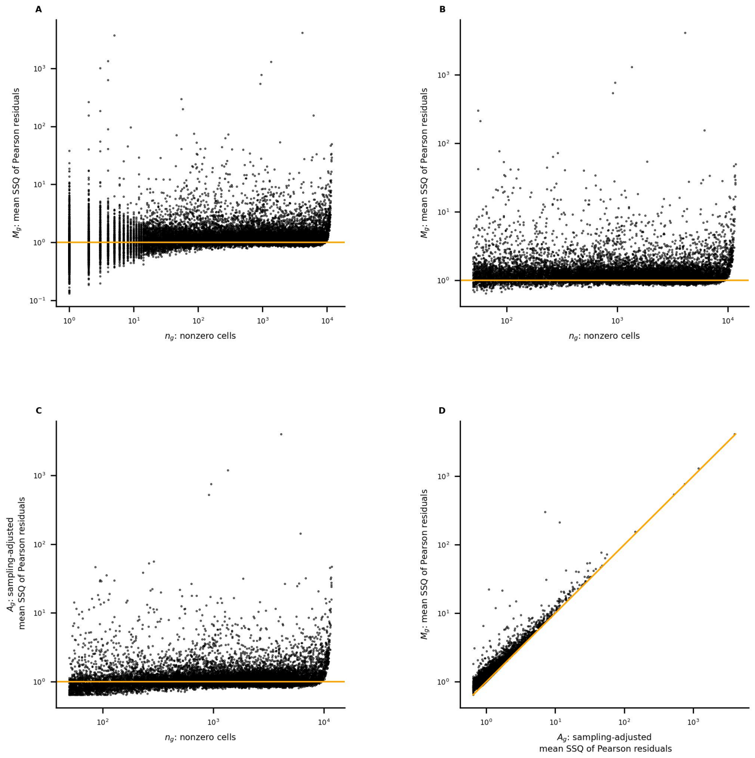


**Figure B-1** 10k brain data: (**A**) the mean SSQ of Pearson residuals $\boldsymbol{M}_g$ vs. the number of nonzero cells $\boldsymbol{n}_g$ for all genes; (**B**) restricting to genes with at least 50 nonzero cells; (**C**) $\boldsymbol{A}_g$ vs. $\boldsymbol{n}_g$; (**D**) the effect of sampling-adjustment: $\boldsymbol{M}_g$ vs. $\boldsymbol{A}_g$

| $\boldsymbol{M}_g$ | $\leq 1$ | $1 < -2$ | $> 2$ | Total |
|---|---|---|---|---|
| $\boldsymbol{n}_g$ | | | | |
| 1-10 | 2,714 | 1,557 | 550 | 4,821 |
| 11-50 | 852 | 1,430 | 187 | 2,469 |
| 51-100 | 217 | 781 | 128 | 1,126 |
| 101-1,000 | 663 | 3,447 | 422 | 4,532 |
| 1,001-10,000 | 1,629 | 5,844 | 483 | 7,956 |
| 10,001+ | 0 | 66 | 110 | 176 |
| Total | 6,075 | 13,125 | 1,880 | 21,080 |

**Table B-1** 10k brain data: relation between $\boldsymbol{n}_g$, the number of nonzero cells, and $\boldsymbol{M}_g$, the mean SSQ of Pearson residuals

| sample $\tilde{S}$<br>sample $S$ | 1-20 | 21-50 | 51-100 | 101-200 | 201-500 | 501-2000 | 2001+ | Total |
|---|---|---|---|---|---|---|---|---|
| 1-20 | 18 | 2 | 0 | 0 | 0 | 0 | 0 | 20 |
| 21-50 | 2 | 24 | 4 | 0 | 0 | 0 | 0 | 30 |
| 51-100 | 0 | 3 | 41 | 6 | 0 | 0 | 0 | 50 |
| 101-200 | 0 | 1 | 5 | 82 | 12 | 0 | 0 | 100 |
| 201-500 | 0 | 0 | 0 | 12 | 253 | 34 | 1 | 300 |
| 501-2000 | 0 | 0 | 0 | 0 | 35 | 1,236 | 229 | 1,500 |
| 2001+ | 0 | 0 | 0 | 0 | 0 | 230 | 10,340 | 10,570 |
| Total | 20 | 30 | 50 | 100 | 300 | 1,500 | 10,570 | 12,570 |

**Table B-2** 10k brain data: comparing ranks of $\boldsymbol{A}_g$ calculated with $S$ and $\tilde{S}$

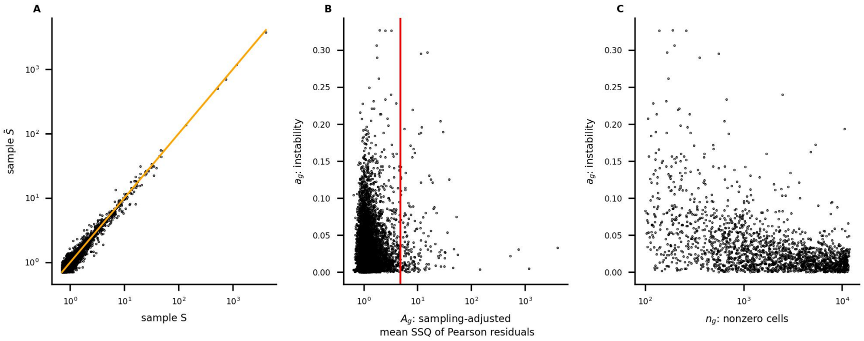


**Figure B-2** 10k brain data: (**A**) large values of $\boldsymbol{A}_g(S)$ and $\boldsymbol{A}_g(\tilde{S})$ agree closely; (**B**) the red vertical line marks the $200^{th}$ ranked value of $\boldsymbol{A}_g$ calculated with all cells; (**C**) for genes with the 2,000 largest values of $\boldsymbol{A}_g$, for which both $\boldsymbol{A}_g(S)$ and $\boldsymbol{A}_g(\tilde{S})$ were also calculated, instability is larger for genes with fewer nonzero cells

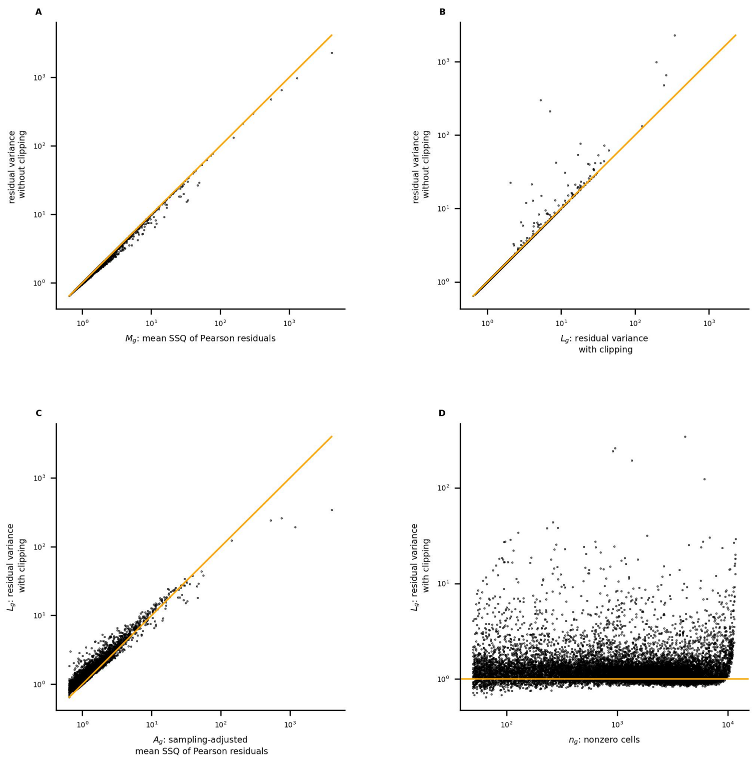


**Figure B-3** 10k brain data: (**A**) scores calculated by ***scanpy*** without clipping agree closely with $\boldsymbol{M}_g$; (**B**) clipping affects scores for very few genes; (**C**) $\boldsymbol{L}_g$ and $\boldsymbol{A}_g$ are large for many of the same genes; (**D**) ***scanpy*** score $\boldsymbol{L}_g$ vs. the number of nonzero cells

| $\boldsymbol{L}_g$ rank<br>$\boldsymbol{A}_g$ rank | 1-20 | 21-50 | 51-100 | 101-200 | 201-500 | 501+ | Total |
|---|---|---|---|---|---|---|---|
| 1-20 | 16 | 4 | 0 | 0 | 0 | 0 | 20 |
| 21-50 | 4 | 21 | 4 | 1 | 0 | 0 | 30 |
| 51-100 | 0 | 5 | 33 | 12 | 0 | 0 | 50 |
| 101-200 | 0 | 0 | 13 | 70 | 17 | 0 | 100 |
| 201-500 | 0 | 0 | 0 | 17 | 220 | 63 | 300 |
| 501+ | 0 | 0 | 0 | 0 | 63 | 13,254 | 13,317 |
| Total | 20 | 30 | 50 | 100 | 300 | 13,317 | 13,817 |

**Table B-3** 10k brain data: comparing ranks of $\boldsymbol{L}_g$ with $\boldsymbol{A}_g$

| sample $\tilde{S}$<br>sample $S$ | 1-20 | 21-50 | 51-100 | 101-200 | 201-500 | 501-2000 | 2001+ | Total |
|---|---|---|---|---|---|---|---|---|
| 1-20 | 18 | 2 | 0 | 0 | 0 | 0 | 0 | 20 |
| 21-50 | 2 | 23 | 5 | 0 | 0 | 0 | 0 | 30 |
| 51-100 | 0 | 5 | 39 | 6 | 0 | 0 | 0 | 50 |
| 101-200 | 0 | 0 | 6 | 83 | 10 | 1 | 0 | 100 |
| 201-500 | 0 | 0 | 0 | 11 | 238 | 50 | 1 | 300 |
| 501-2000 | 0 | 0 | 0 | 0 | 52 | 1,171 | 277 | 1,500 |
| 2001+ | 0 | 0 | 0 | 0 | 0 | 278 | 10,292 | 10,570 |
| Total | 20 | 30 | 50 | 100 | 300 | 1,500 | 10,570 | 12,570 |

**Table B-4** 10k brain data: comparing ranks of $\boldsymbol{L}_g$ calculated with $S$ and $\tilde{S}$

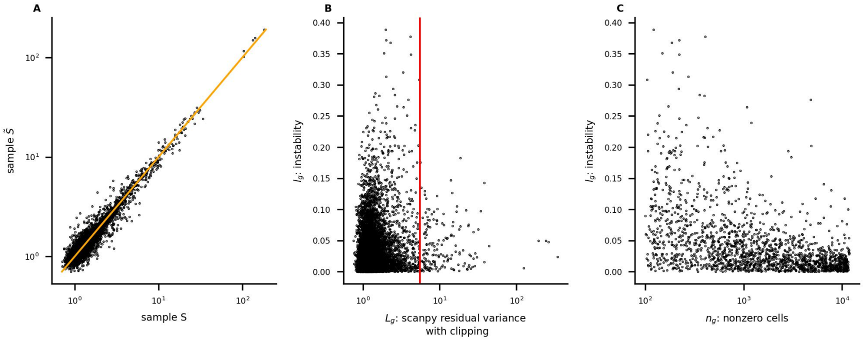


**Figure B-4** 10k brain data: (**A**) large values of $\boldsymbol{L}_g(S)$ and $\boldsymbol{L}_g(\tilde{S})$ agree closely; (**B**) the red vertical line marks the $200^{th}$ ranked value of $\boldsymbol{L}_g$ calculated with all cells; (**C**) for genes with the 2,000 largest values of $\boldsymbol{L}_g$, for which both $\boldsymbol{L}_g(S)$ and $\boldsymbol{L}_g(\tilde{S})$ were also calculated, instability is larger for genes with fewer nonzero cells

The distributions of $\boldsymbol{a}_g$ and $\boldsymbol{l}_g$ were compared by performing Mann-Whitney and Kolmogorov-Smirnov tests (restricting to genes represented in Figures B-2C and B-4C).

The Mann-Whitney one-sided test rejects the null hypothesis that $\boldsymbol{l}_g$ and $\boldsymbol{a}_g$ have the same distribution in favor of the alternative that the distribution of $\boldsymbol{l}_g$ is stochastically greater than that of $\boldsymbol{a}_g$ (p=3e-5). Similarly, the Kolmogorov-Smirnov one-sided test rejects the null hypothesis in favor of the alternative that the CDF of $\boldsymbol{l}_g$ is less than that of $\boldsymbol{a}_g$ (p=2e-6).

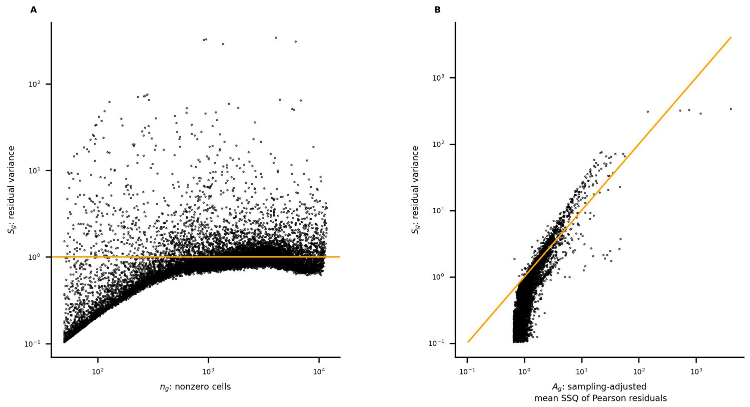


**Figure B-5** 10k brain data: (**A**) ***SCTransform*** assigns very low scores to most genes with few nonzero cells; (**B**) $\boldsymbol{S}_g$ and $\boldsymbol{A}_g$ are large for many of the same genes

| $\boldsymbol{S}_g$ rank / $\boldsymbol{A}_g$ rank | 1-20 | 21-50 | 51-100 | 101-200 | 201-500 | 501+ | Total |
|---|---|---|---|---|---|---|---|
| 1-20 | 11 | 3 | 2 | 0 | 2 | 2 | 20 |
| 21-50 | 9 | 12 | 2 | 2 | 1 | 4 | 30 |
| 51-100 | 0 | 15 | 22 | 7 | 4 | 2 | 50 |
| 101-200 | 0 | 0 | 24 | 48 | 19 | 9 | 100 |
| 201-500 | 0 | 0 | 0 | 42 | 182 | 76 | 300 |
| 501+ | 0 | 0 | 0 | 1 | 92 | 13,224 | 13,317 |
| Total | 20 | 30 | 50 | 100 | 300 | 13,317 | 13,817 |

**Table B-5** 10k brain data: comparing ranks of $\boldsymbol{S}_g$ with $\boldsymbol{A}_g$

| sample $\tilde{S}$ / sample $S$ | 1-20 | 21-50 | 51-100 | 101-200 | 201-500 | 501-2000 | 2001+ | Total |
|---|---|---|---|---|---|---|---|---|
| 1-20 | 15 | 5 | 0 | 0 | 0 | 0 | 0 | 20 |
| 21-50 | 5 | 19 | 6 | 0 | 0 | 0 | 0 | 30 |
| 51-100 | 0 | 6 | 36 | 7 | 1 | 0 | 0 | 50 |
| 101-200 | 0 | 0 | 8 | 77 | 15 | 0 | 0 | 100 |
| 201-500 | 0 | 0 | 0 | 13 | 242 | 45 | 0 | 300 |
| 501-2000 | 0 | 0 | 0 | 3 | 41 | 1,249 | 207 | 1,500 |
| 2001+ | 0 | 0 | 0 | 0 | 1 | 206 | 10,363 | 10,570 |
| Total | 20 | 30 | 50 | 100 | 300 | 1,500 | 10,570 | 12,570 |

**Table B-6** 10k brain data: comparing ranks of $\boldsymbol{S}_g$ calculated with $S$ and $\tilde{S}$

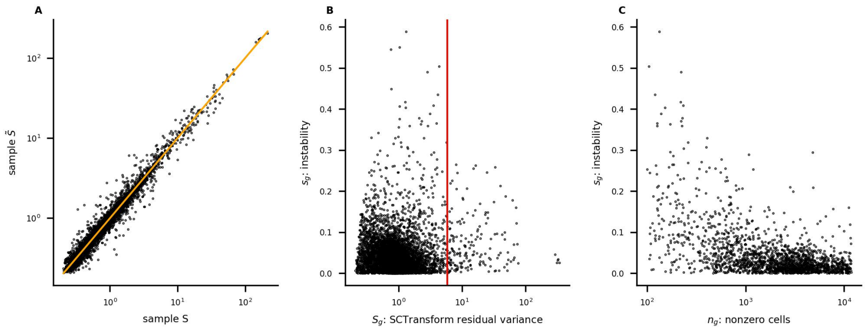


**Figure B-6** 10k brain data: (**A**) large values of $\boldsymbol{S}_g(S)$ and $\boldsymbol{S}_g(\tilde{S})$ agree closely; (**B**) the red vertical line marks the $200^{th}$ ranked value of $\boldsymbol{S}_g$ calculated with all cells; (**C**) for genes with the 2,000 largest values of $\boldsymbol{S}_g$, for which both $\boldsymbol{S}_g(S)$ and $\boldsymbol{S}_g(\tilde{S})$ were also calculated, instability is larger for genes with fewer nonzero cells

The distributions of $\boldsymbol{a}_g$ and $\boldsymbol{s}_g$ were compared by performing Mann-Whitney and Kolmogorov-Smirnov tests (restricting to genes represented in Figures B-2C and B-6C).

The Mann-Whitney one-sided test rejects the null hypothesis that $\boldsymbol{s}_g$ and $\boldsymbol{a}_g$ have the same distribution in favor of the alternative that the distribution of $\boldsymbol{s}_g$ is stochastically greater than that of $\boldsymbol{a}_g$ (p=2e-7). Similarly, the Kolmogorov-Smirnov one-sided test rejects the null hypothesis in favor of the alternative that the CDF of $\boldsymbol{s}_g$ is less than that of $\boldsymbol{a}_g$ (p=2e-5).

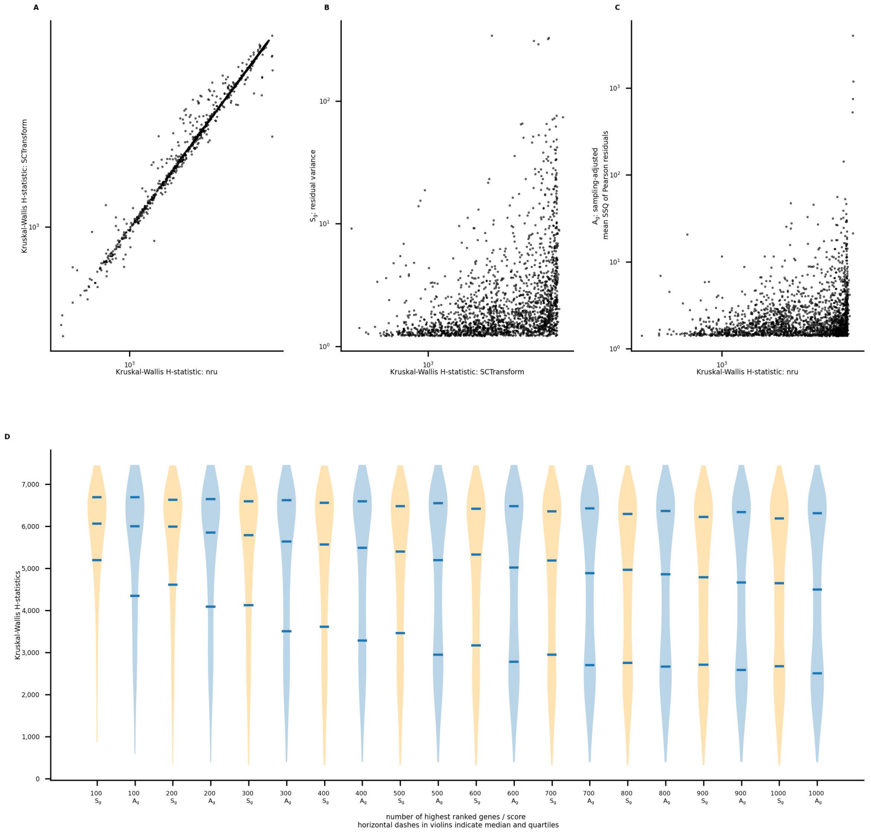


**Figure B-7** 10k brain data: Kruskal-Wallis H-statistics were calculated using a 10-cluster segmentation; (**A**) for genes ranked in the top 2,000 by both methods, H-statistics are strongly correlated

Using Pearson residuals calculated with ***nru*** and ***SCTransform***, Kruskal-Wallis tests were performed for genes with the 2,000 largest values of $\boldsymbol{A}_g$ and $\boldsymbol{S}_g$.

The violin plots (B-7D) compare the distributions of H-statistics for residuals calculated with ***nru*** and ***SCTransform*** for the corresponding method's highest ranked $100, \cdots, 1,000$ genes.

For the sets of 500, 600, and 700 genes, one-sided Kolmogorov-Smirnov tests reject the null hypothesis in favor of the alternative that the CDF of H-statistics for ***nru*** is greater than the CDF of the H-statistics for ***SCTransform*** ($p < 0.05$). This analysis provides no reason to prefer one method over the other.

| genes | $\boldsymbol{S}_g$ | $\boldsymbol{A}_g$ |
|---|---|---|
| 50 | 0.21 | 0.41 |
| 100 | -0.04 | 0.07 |
| 200 | 0.10 | 0.15 |
| 500 | 0.27 | 0.30 |
| 1000 | 0.31 | 0.21 |
| 2000 | 0.47 | 0.26 |

**Table B-7** 10k brain data: Spearman correlations between H-statistics and $\boldsymbol{S}_g$ or $\boldsymbol{A}_g$ for genes with the largest 50, 100, 200, 500, 1,000, and 2,000 H-statistics

For the sets of genes with the largest 50, 100, 200, 500, 1,000, and 2,000 H-statistics, Spearman correlations between H-statistics and $\boldsymbol{S}_g$ or $\boldsymbol{A}_g$ were calculated. They are summarized in Table B-7. For the sets of 50 genes with the largest H-statistics, correlations between $\boldsymbol{A}_g$ and H-statistics calculated with ***nru*** are larger; for the sets of 2,000 genes, correlations between $\boldsymbol{S}_g$ and H-statistics calculated with ***SCTransform*** are larger.

### 10k heart data

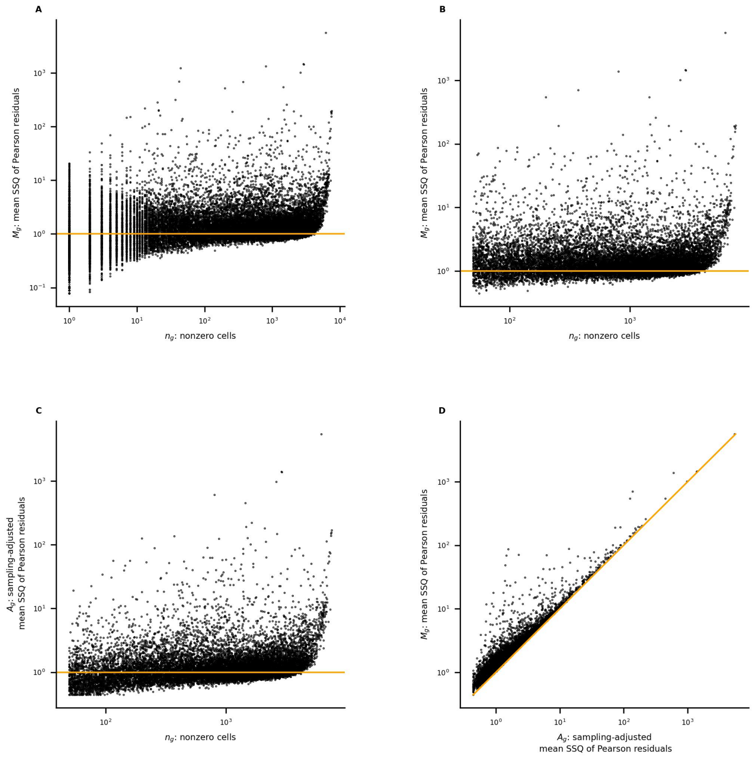


**Figure H-1** 10k heart data: (**A**) the mean SSQ of Pearson residuals $\boldsymbol{M}_g$ vs. the number of nonzero cells $\boldsymbol{n}_g$ for all genes; (**B**) restricting to genes with at least 50 nonzero cells; (**C**) $\boldsymbol{A}_g$ vs. $\boldsymbol{n}_g$; (**D**) the effect of sampling-adjustment: $\boldsymbol{M}_g$ vs. $\boldsymbol{A}_g$

| $\boldsymbol{M}_g$ <br> $\boldsymbol{n}_g$ | $\leq 1$ | $1 < -2$ | $> 2$ | Total |
|---|---|---|---|---|
| 1-10 | 3,037 | 1,115 | 1,048 | 5,200 |
| 11-50 | 1,061 | 1,195 | 737 | 2,993 |
| 51-100 | 438 | 575 | 291 | 1,304 |
| 101-1,000 | 1,598 | 3,523 | 1,260 | 6,381 |
| 1,001-10,000 | 1,240 | 4,160 | 1,343 | 6,743 |
| 10,001+ | 0 | 0 | 0 | 0 |
| Total | 7,374 | 10,568 | 4,679 | 22,621 |

**Table H-1** 10k heart data: relation between $\boldsymbol{n}_g$, the number of nonzero cells, and $\boldsymbol{M}_g$, the mean SSQ of Pearson residuals

| sample $\tilde{S}$<br>sample $S$ | 1-20 | 21-50 | 51-100 | 101-200 | 201-500 | 501-2000 | 2001+ | Total |
|---|---|---|---|---|---|---|---|---|
| 1-20 | 18 | 1 | 0 | 0 | 0 | 1 | 0 | 20 |
| 21-50 | 2 | 25 | 2 | 0 | 0 | 0 | 1 | 30 |
| 51-100 | 0 | 4 | 42 | 3 | 0 | 0 | 1 | 50 |
| 101-200 | 0 | 0 | 6 | 81 | 11 | 2 | 0 | 100 |
| 201-500 | 0 | 0 | 0 | 16 | 262 | 22 | 0 | 300 |
| 501-2000 | 0 | 0 | 0 | 0 | 27 | 1,306 | 167 | 1,500 |
| 2001+ | 0 | 0 | 0 | 0 | 0 | 169 | 10,826 | 10,995 |
| Total | 20 | 30 | 50 | 100 | 300 | 1,500 | 10,995 | 12,995 |

**Table H-2** 10k heart data: comparing ranks of $\boldsymbol{A}_g$ calculated with $S$ and $\tilde{S}$

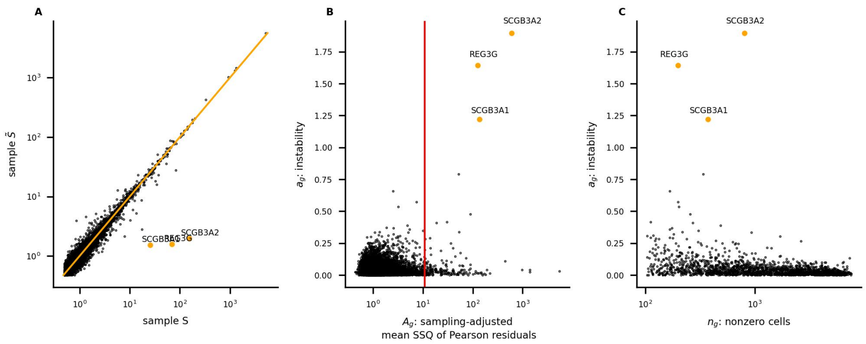


**Figure H-2** 10k heart data: (**A**) large values of $\boldsymbol{A}_g(S)$ and $\boldsymbol{A}_g(\tilde{S})$ agree closely except for the three outliers REG3G, SCGB3A1, and SCGB3A2; (**B**) the red vertical line marks the $200^{th}$ ranked value of $\boldsymbol{A}_g$ calculated with all cells; (**C**) for genes with the 2,000 largest values of $\boldsymbol{A}_g$, for which both $\boldsymbol{A}_g(S)$ and $\boldsymbol{A}_g(\tilde{S})$ were also calculated, instability is larger for genes with fewer nonzero cells

The large differences between $\boldsymbol{A}_g(S)$ and $\boldsymbol{A}_g(\tilde{S})$ for the genes REG3G, SCGB3A1, and SCGB3A2 may indicate a quality control issue – counts concentrated on a small number of cells:

- REG3G: 1 cell is responsible for 26% of its total count, 2 cells for 49%, 7 for 94%
- SCGB3A1: 1 cell is responsible for 23% of its total count, 2 cells for 45%, 6 for 93%
- SCGB3A2: 1 cell is responsible for 27% of its total count, 2 cells for 42%, 7 for 96%

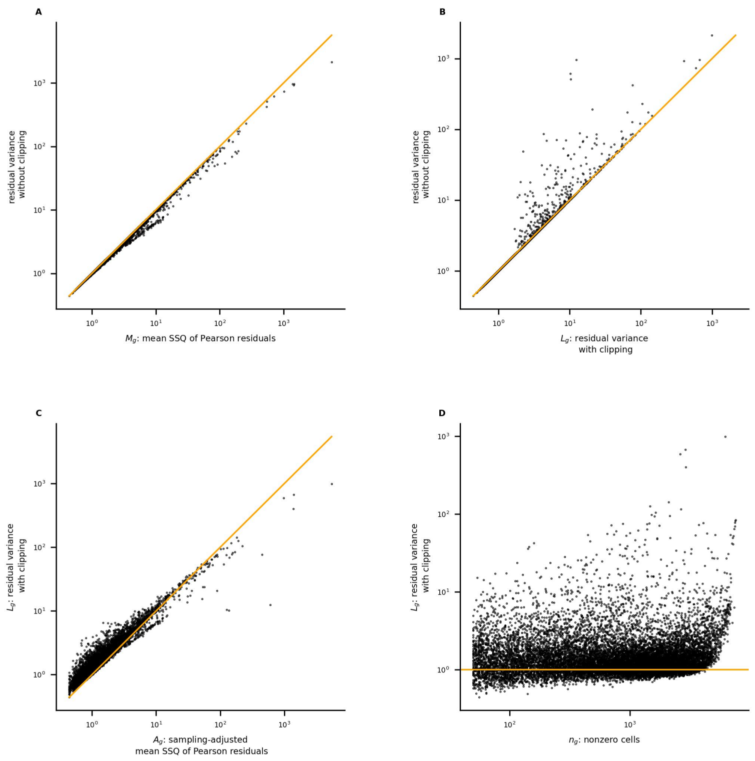


**Figure H-3** 10k heart data: (**A**) scores calculated by ***scanpy*** without clipping agree closely with $\boldsymbol{M}_g$; (**B**) clipping affects scores for very few genes; (**C**) $\boldsymbol{L}_g$ and $\boldsymbol{A}_g$ are large for many of the same genes; (**D**) ***scanpy*** score $\boldsymbol{L}_g$ vs. the number of nonzero cells

| $\boldsymbol{L}_g$ rank<br>$\boldsymbol{A}_g$ rank | 1-20 | 21-50 | 51-100 | 101-200 | 201-500 | 501+ | Total |
|---|---|---|---|---|---|---|---|
| 1-20 | 15 | 2 | 0 | 1 | 2 | 0 | 20 |
| 21-50 | 5 | 21 | 2 | 2 | 0 | 0 | 30 |
| 51-100 | 0 | 7 | 39 | 4 | 0 | 0 | 50 |
| 101-200 | 0 | 0 | 9 | 75 | 16 | 0 | 100 |
| 201-500 | 0 | 0 | 0 | 18 | 219 | 63 | 300 |
| 501+ | 0 | 0 | 0 | 0 | 63 | 13,909 | 13,972 |
| Total | 20 | 30 | 50 | 100 | 300 | 13,972 | 14,472 |

**Table H-3** 10k heart data: comparing ranks of $\boldsymbol{L}_g$ with $\boldsymbol{A}_g$

| sample $\tilde{S}$ <br> sample $S$ | 1-20 | 21-50 | 51-100 | 101-200 | 201-500 | 501-2000 | 2001+ | Total |
|---|---|---|---|---|---|---|---|---|
| 1-20 | 19 | 1 | 0 | 0 | 0 | 0 | 0 | 20 |
| 21-50 | 1 | 29 | 0 | 0 | 0 | 0 | 0 | 30 |
| 51-100 | 0 | 0 | 47 | 3 | 0 | 0 | 0 | 50 |
| 101-200 | 0 | 0 | 3 | 87 | 10 | 0 | 0 | 100 |
| 201-500 | 0 | 0 | 0 | 10 | 251 | 39 | 0 | 300 |
| 501-2000 | 0 | 0 | 0 | 0 | 38 | 1,253 | 209 | 1,500 |
| 2001+ | 0 | 0 | 0 | 0 | 1 | 208 | 10,786 | 10,995 |
| Total | 20 | 30 | 50 | 100 | 300 | 1,500 | 10,995 | 12,995 |

**Table H-4** 10k heart data: comparing ranks of $\boldsymbol{L}_g$ calculated with $S$ and $\tilde{S}$

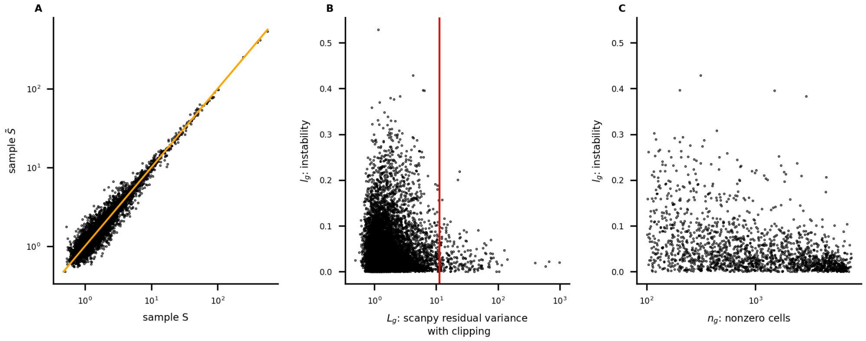


**Figure H-4** 10k heart data: (**A**) large values of $\boldsymbol{L}_g(S)$ and $\boldsymbol{L}_g(\tilde{S})$ agree closely; (**B**) the red vertical line marks the $200^{th}$ ranked value of $\boldsymbol{L}_g$ calculated with all cells; (**C**) for genes with the 2,000 largest values of $\boldsymbol{L}_g$, for which both $\boldsymbol{L}_g(S)$ and $\boldsymbol{L}_g(\tilde{S})$ were also calculated, instability is larger for genes with fewer nonzero cells

The distributions of $\boldsymbol{a}_g$ and $\boldsymbol{l}_g$ were compared by performing Mann-Whitney and Kolmogorov-Smirnov tests (restricting to genes represented in Figures H-2C and H-4C).

The Mann-Whitney one-sided test rejects the null hypothesis that $\boldsymbol{l}_g$ and $\boldsymbol{a}_g$ have the same distribution in favor of the alternative that the distribution of $\boldsymbol{l}_g$ is stochastically greater than that of $\boldsymbol{a}_g$ (p=0.001). Similarly, the Kolmogorov-Smirnov one-sided test rejects the null hypothesis in favor of the alternative that the CDF of $\boldsymbol{l}_g$ is less than that of $\boldsymbol{a}_g$ (p=0.002).

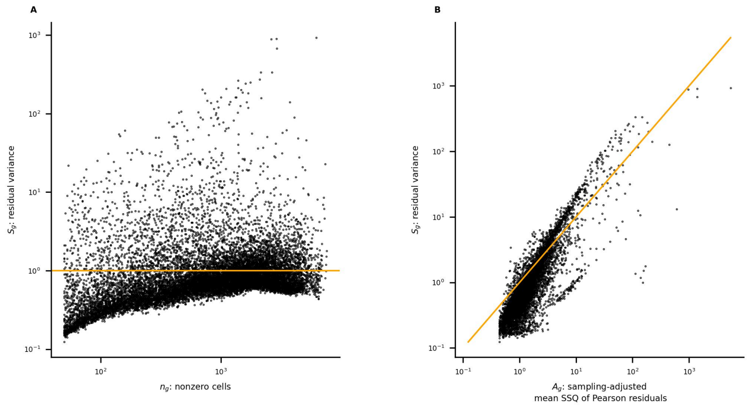


**Figure H-5** 10k heart data: (**A**) ***SCTransform*** assigns very low scores to most genes with few nonzero cells; (**B**) $\boldsymbol{S_g}$ and $\boldsymbol{A_g}$ are large for many of the same genes

| $\boldsymbol{S_g}$ rank<br>$\boldsymbol{A_g}$ rank | 1-20 | 21-50 | 51-100 | 101-200 | 201-500 | 501+ | Total |
|---|---|---|---|---|---|---|---|
| 1-20 | 9 | 3 | 0 | 0 | 3 | 5 | 20 |
| 21-50 | 9 | 8 | 7 | 2 | 3 | 1 | 30 |
| 51-100 | 2 | 18 | 19 | 6 | 3 | 2 | 50 |
| 101-200 | 0 | 1 | 24 | 46 | 6 | 23 | 100 |
| 201-500 | 0 | 0 | 0 | 46 | 140 | 114 | 300 |
| 501+ | 0 | 0 | 0 | 0 | 145 | 13,827 | 13,972 |
| Total | 20 | 30 | 50 | 100 | 300 | 13,972 | 14,472 |

**Table H-5** 10k heart data: comparing ranks of $\boldsymbol{S_g}$ with $\boldsymbol{A_g}$

| sample $\tilde{S}$<br>sample $S$ | 1-20 | 21-50 | 51-100 | 101-200 | 201-500 | 501-2000 | 2001+ | Total |
|---|---|---|---|---|---|---|---|---|
| 1-20 | 18 | 2 | 0 | 0 | 0 | 0 | 0 | 20 |
| 21-50 | 2 | 25 | 3 | 0 | 0 | 0 | 0 | 30 |
| 51-100 | 0 | 3 | 42 | 5 | 0 | 0 | 0 | 50 |
| 101-200 | 0 | 0 | 5 | 87 | 8 | 0 | 0 | 100 |
| 201-500 | 0 | 0 | 0 | 8 | 243 | 49 | 0 | 300 |
| 501-2000 | 0 | 0 | 0 | 0 | 49 | 1,229 | 222 | 1,500 |
| 2001+ | 0 | 0 | 0 | 0 | 0 | 222 | 10,773 | 10,995 |
| Total | 20 | 30 | 50 | 100 | 300 | 1,500 | 10,995 | 12,995 |

**Table H-6** 10k heart data: comparing ranks of $\boldsymbol{S_g}$ calculated with $S$ and $\tilde{S}$

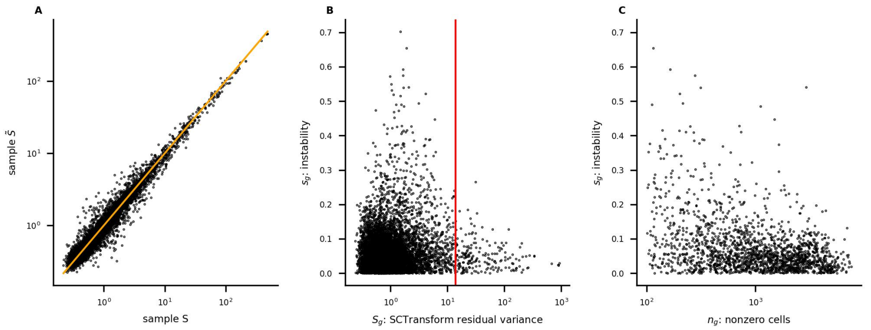


**Figure H-6** 10k heart data: (**A**) large values of $\boldsymbol{S}_g(S)$ and $\boldsymbol{S}_g(\tilde{S})$ agree closely; (**B**) the red vertical line marks the $200^{th}$ ranked value of $\boldsymbol{S}_g$ calculated with all cells; (**C**) for genes with the 2,000 largest values of $\boldsymbol{S}_g$, for which both $\boldsymbol{S}_g(S)$ and $\boldsymbol{S}_g(\tilde{S})$ were also calculated, instability is larger for genes with fewer nonzero cells

The distributions of $\boldsymbol{a}_g$ and $\boldsymbol{s}_g$ were compared by performing Mann-Whitney and Kolmogorov-Smirnov tests (restricting to genes represented in Figures H-2C and H-6C).

The Mann-Whitney one-sided test rejects the null hypothesis that $\boldsymbol{s}_g$ and $\boldsymbol{a}_g$ have the same distribution in favor of the alternative that the distribution of $\boldsymbol{s}_g$ is stochastically greater than that of $\boldsymbol{a}_g$ (p=0). Similarly, the Kolmogorov-Smirnov one-sided test rejects the null hypothesis in favor of the alternative that the CDF of $\boldsymbol{s}_g$ is less than that of $\boldsymbol{a}_g$ (p=0).

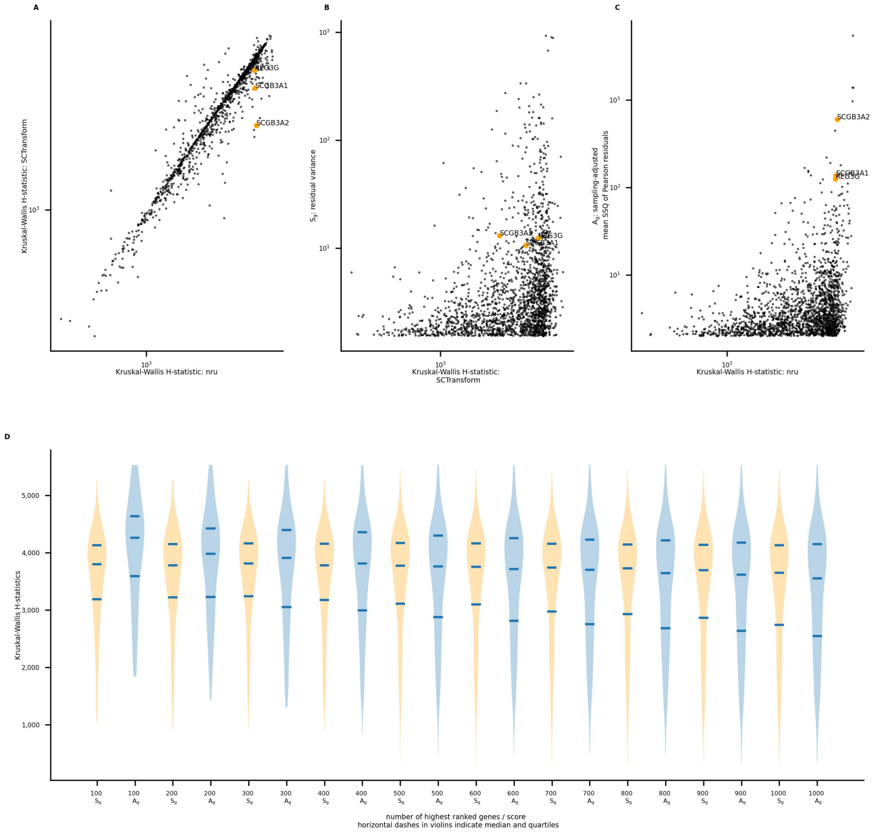


**Figure H-7** 10k heart data: Kruskal-Wallis H-statistics were calculated using a 10-cluster segmentation; (**A**) for genes ranked in the top 2,000 by both methods, H-statistics are strongly correlated; (**D**) Mann-Whitney and Kolmogorov-Smirnov tests find the H-statistics for ***nru*** to be larger for the top ranked $100, \cdots, 400$ genes

The three outlier genes identified in Figure H-2 (REG3G, SCGB3A1, and SCGB3A2) are notated. They are ranked among the top 20 by ***nru*** and in the range 210 to 260 by ***SCTransform***.

The violin plots (H-7D) compare the distributions of H-statistics for residuals calculated with ***nru*** and ***SCTransform*** for the corresponding method's highest ranked $100, \cdots, 1,000$ genes.

For the sets of 400 genes or fewer, the one-sided Mann-Whitney test rejects the null hypothesis that the distributions of H-statistics for ***nru*** and ***SCTransform*** are equal, in favor of the alternative that the distribution for ***nru*** is stochastically greater ($p < 0.04$). For the same sets of genes, Kolmogorov-Smirnov one-sided tests reject the null hypothesis in favor of the alternative that the CDF of H-statistics for ***nru*** is less than the CDF of the H-statistics for ***SCTransform*** ($p < 0.002$). We note that for the sets of 800 genes or more, one-sided KS tests *also* reject the null hypothesis in favor of the alternative that the CDF of H-statistics for ***nru*** is *greater* than the CDF of the H-statistics for ***SCTransform*** ($p < 0.01$).

| genes | $\boldsymbol{S}_g$ | $\boldsymbol{A}_g$ |
|---|---|---|
| 50 | -0.02 | 0.57 |
| 100 | 0.01 | 0.35 |
| 200 | -0.08 | 0.19 |
| 500 | 0.04 | 0.26 |
| 1000 | 0.10 | 0.17 |
| 2000 | 0.38 | 0.33 |

**Table H-7** 10k heart data: Spearman correlations between H-statistics and $\boldsymbol{S}_g$ or $\boldsymbol{A}_g$ for genes with the largest 50, 100, 200, 500, 1,000, and 2,000 H-statistics

Correlations between H-statistics and $\boldsymbol{S}_g$ or $\boldsymbol{A}_g$ were calculated for varying numbers of genes with the highest scores. For the sets of 500 genes or fewer, correlations between $\boldsymbol{A}_g$ and H-statistics calculated with $\boldsymbol{nru}$ are larger.
